# Cell-of-origin epigenome underlies SS18::SSX-mediated transformation

**DOI:** 10.1101/2024.05.15.594021

**Authors:** Lesley A. Hill, R. Wilder Scott, Lauren Martin, Martin Arostegui, George Davenport, Marcos Vemon, Jakob Hofvander, Xue Qi Wang, Jinxiu Li, Torsten O. Nielsen, Kevin B. Jones, Martin Hirst, T. Michael Underhill

## Abstract

Synovial sarcoma is an aggressive soft-tissue malignancy that is characterized by a pathognomonic t(X;18)(p11.2;q11.2) translocation, which produces the fusion oncogene named *SS18::SSX*. Despite recent advancements in our understanding of synovial sarcoma biology, the cell-of-origin remains undefined. A mesenchymal stromal cell (MSC) specific CreERT2 line was employed to express *SS18::SSX* in fibroblasts and related cell types, resulting in 100% penetrant synovial sarcoma development in mice, with a median latency period of 16.2 ± 2.5 weeks. Murine tumours exhibited high concordance with human synovial sarcoma sub-types at the histological and molecular levels^1^. Genetic refinement of the cell-of-origin revealed that synovial sarcomas derive from a rare *Hic1*^+^ *Pdgfra*^+^ *Lgr5*^+^ fibroblastic population. Furthermore, longitudinal multi-omic profiling along the transformation continuum revealed the step-wise acquisition of a transformed phenotype initiated by the loss of a mature fibroblastic profile and subsequently, the gradual unmasking of an epigenetically embedded embryonic MSC program. Adult and embryonic MSCs exhibited overlapping H2AK119ub and H3K4me3/H3K27me3 (bivalent) histone marks, while SS18::SSX-mediated transformation culminated in the widespread loss of H3K27me3 at these genes and their consequent transcription. Collectively, these studies define a rare MSC context, conducive for SS18::SSX-mediated transformation, and demonstrate that SS tumorigenesis involves the induction and maintenance of an embryonic-like MSC phenotype.

---

Synovial sarcoma (SyS) is a malignant soft-tissue sarcoma that primarily affects adolescents or young adults, and can arise in virtually any area of the body. The biologic behaviour of SyS can vary depending on a variety of factors, including histologic subtype, tumour size, and age at diagnosis. The 5- and 10-year survival rates have been estimated at approximately 60% and 50%, respectively^2,3^. SySs typically arise in the deep soft tissues of the extremities, often near (but outside) large joints^4^, although truncal^3^, as well as head and neck^4^ sites are not uncommon.

SyS is characterized by a t(X;18)(p11.2;q11.2) translocation that produces one of three fusion oncogenes: *SS18::SSX1, SS18::SSX2,* or less commonly, *SS18::SSX4*^5–8^. While the SS18::SSX fusion oncoprotein is both necessary and sufficient for synovial sarcomagenesis, it is unclear exactly how this protein causes neoplastic transformation in cells expressing the oncogene. The native SS18 protein is a core subunit of BAF complexes, including CBAF and GBAF, and the presence of the SS18::SSX fusion protein was shown to disrupt BAF function, leading to altered transcription, genome-wide^9–11^. One of the main roles of BAF complexes is to oppose Polycomb Repressive Complexes (PRC1/2). PRC complexes are histone modifiers that facilitate transcriptional repression through the placement of specific histone marks, such as trimethylation of lysine 27 in histone H3 (H3K27me3) and monoubiquitination of lysine 119 in histone H2A (H2AK119ub1). They are also responsible for maintaining transcriptional repression of “bivalently marked” promoters – those bearing both H3K27me3 and trimethylation of lysine 4 in histone H3 (H3K4me3) marks – which leave genes poised for expression upon the loss of H3K27me3 repression^12–14^. In addition to its interaction with BAF complexes, SS18::SSX has been shown to associate with PRCs, specifically serving as a bridge between PRC1.1 and BAF complexes^15^. Importantly, the C-terminus of SSX has been demonstrated to interact with H2AK119ub-rich regions, targeting SS18::SSX to these loci^10,16,17^. SS18::SSX also alters the stoichiometry of the defined BAF complexes - PBAF, GBAF and CBAF - with the latter being preferentially degraded^11^. While the exact molecular mechanisms driving SyS remain unknown, there is clear evidence for epigenetic dysregulation through interactions with activating BAF complexes and repressive PRC complexes^18^. The SS18::SSX fusion protein appears to cause widespread transcriptional dysregulation, associated with both gene activation and repression, leading to a proliferative phenotype and a neural/developmental expression pattern that is / among soft-tissue sarcomas^19–24^.

The cell-of-origin for many sarcomas is thought to be mesenchymal in nature, primarily due to the tissue compartments within which they typically arise. Accordingly, the discrete identification of a specific cell with strong origination potential would provide a valuable normal comparator to better understand SS18::SSX-mediated transformation. To address this, we used a recently generated CreERT2 mouse line, *Hic1^CreERT2^*, that is restricted to mesenchymal stromal cells (MSCs) throughout adult tissues, including fibroblasts and pericytes, to target human *SS18::SSX2* expression to this cell population^25–27^. Mice presented, at 100% penetrance, with tumours that were characteristic of human SyS. These developed at a consistent latency and appeared at select anatomical sites, indicating that Hic1^+^ MSCs represent a potential cell of origin for SyS. ScRNA-seq trajectory analysis along the transformation continuum revealed parallels to molecular events that are observed when reprogramming fibroblasts for iPSC generation, with the subsequent emergence of an embryonic transcriptional signature^28,29^. Consistent with this, transcriptomic and epigenomic analyses revealed that the neural/developmental transcriptional signature, characteristic of SyS^22^, was embedded in the MSC epigenome, as a constellation of histone marks consistent with bivalency. This included H3K4me3, H3K27me3 and H2AK119ub1, with the latter serving as a targeting moiety for SS18::SSX. Embryonic Hic1^+^ MSCs, characterized shortly after their emergence, exhibited a similar pattern of bivalently marked regions, indicating that the adult MSC epigenome is a by-product of their embryonic heritage. Collectively, these findings define cellular and epigenomic features conducive for SS18::SSX-mediated transformation, and the consequent emergence of a unique transcriptional SyS landscape, reflective of this deadly cancer.

## A cell-of-origin for synovial sarcoma

To directly test if SyS arise from *Hic1*^+^ MSCs, a *Hic1^CreERT2^* line, was bred with a conditional allele of *SS18::SSX2*, also containing enhanced green fluorescent protein on an internal ribosomal entry site (IRES-EGFP) cassette (*Rosa^hSS2^*). *SS18::SSX2* expression was induced with tamoxifen (TAM) at ∼8 weeks of age (Fig. 1a and 1b). Mice heterozygous for the hSS2 allele presented with tumours at a median latency of ∼ 66 weeks (post-TAM; Fig. 1c), and these appeared at various anatomical sites (i.e., trunk, craniofacial, and appendicular regions), reflective of previous studies^30,31^. Earlier reports also showed that homozygosity of the hSS2 allele shortened tumour latency^11^. In line with these findings, the tumour latency of mice homozygous for the hSS2 allele in the *Hic^CreERT2^* background (herein referred to as the *Hic1; hSS2* mice) was shortened to 16.2 ± 2.5 weeks (Fig. 1c), with females exhibiting an ∼2-week shorter latency in comparison to males (Extended Data Fig. 1a). In contrast to the restricted expression of *SS18::SSX* using *Hic1^CreERT2^*, near ubiquitous expression of *SS18::SSX2* in adult mice, using a UBC-CreERT2 line, led to clinical end-point within three days following TAM administration (Extended Data Fig. 1b). GFP^+^ tumours (*hSS2* allele, IRES-EGFP cassette expression) appeared in the tongue (tSRC) with 100% penetrance (Fig. 1d), in the stifle area (sSRC; Fig. 1d) and less frequently at other sites (Extended Data Fig. 1c). Histological analysis of tongue and stifle tumours revealed characteristic features of monophasic (Mp; i.e., spindle to oval shaped cells with evenly dispersed chromatin and inconspicuous nucleoli) and poorly differentiated (PD; i.e., increased cellularity, ovoid cells with prominent nucleoli) SyS, and positive staining for SS18::SSX (Fig. 1e). In the tongue, Mp and PD SyS appeared at similar frequency, with islands of biphasic (Bi) tumour containing glandular-like structures detected at a lower incidence (Extended Fig. 1d). Consistent with human SyS, epithelial membrane antigen 1 (EMA1 or MUC1) was detected in a diffuse pattern in Mp tumours or on the apical surface of tumour-associated pseudoglandular structures (Extended Data Fig. 1d and e). Tumours broadly expressed nuclear TLE, a marker commonly used to differentiate SyS from other sarcomas^32^, while surrounding margins (GFP**^-^** and SS18::SSX**^-^**) exhibited punctate TLE expression (Extended Data Fig. 1f, g). In a variety of tissues, *Hic1* identifies quiescent MSCs^25–27^, and congruent with this, under homeostasis, 0.5% of tdTomato(tdTom)-labelled (*Hic1^CreERT2^*; *Rosa^tdTom/+^; herein referred to as Hic1; tdTom* mice) tongue MSCs exhibited EdU incorporation after 5 days of EdU treatment (Extended Data Fig. 1h and i). In contrast, neoplastic GFP^+^ cells from end-point tongue tumours exhibited a significant increase in EdU incorporation (20.8%; p < 0.0001%, compared to control, n=3) (Extended Fig 1h-k). Furthermore, SyS exhibits a unique neural/developmental transcriptional signature^22^, and both tongue and stifle tumours (in comparison to corresponding control regions) exhibited marked up-regulation of numerous orthologous transcripts (i.e., *Cdx2, Lhx2, Nkx6-2, Mnx1, Pax2, Pax6, Pou3f3, Sox1, Sox3, Uncx*) associated with this profile (Fig. 1g and Supplementary Table 1). Euclidian distance and principal component analysis revealed high concordance between replicates, with ∼85% of the variance due to anatomical site (Extended Data Fig. 1l, m). Collectively, the histological and molecular findings are congruent with the generation of SyS tumours from a *Hic1*^+^ MSC.

**Fig. 1:**
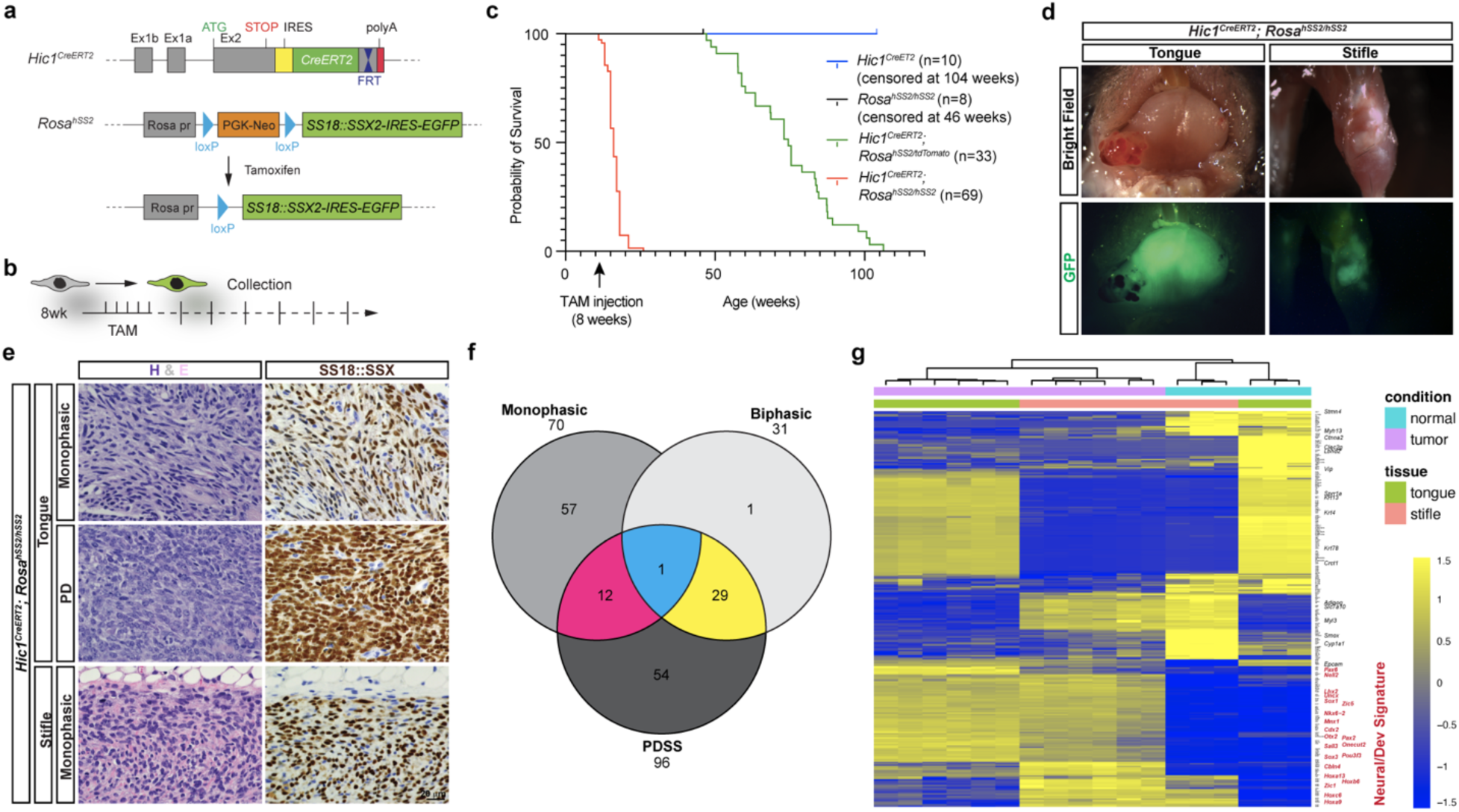
Induction of *SS18::SSX* in *Hic1*^+^ MSCs leads to SyS formation. **a,** Schematic overview of the genetic strategy used to conditionally express *SS18::SSX* in *Hic1*^+^ MSCs using *Hic1^CreERT2^* and *Rosa^hSS2^* alleles (conditional *SS18::SSX2*-IRES-EGFP). **b**, Overview of the experimental plan, whereby *SS18::SSX* was induced at 8 weeks of age, followed by sample collection at various time points. **c**, Kaplan-Meier survival curves following induction of *SS18::SSX2* expression; animal numbers for each genetic background are indicated in parentheses. **d**, Whole-mount images of GFP^+^ tumours at the indicated sites. **e**, Representative images of tumour sub-types following H&E staining and anti-SS18::SSX IHC. **f**, Venn diagram of tumour sub-type abundance from analysis of H&E histological samples across tongue and stifle tumours. **g**, Heatmap showing differential gene expression (top 50 up and down regulated transcripts per group) between tumour samples, and controls collected from the tumour origin site (normal tongue and stifle region).

In skeletal muscle (SkM), heart, and skin, *Hic1* identifies 2 major mesenchymal stromal cell constituents, fibroblasts (Fb; *Pdgfra*^+^, *Dpt*^+^, *Col15a1*^+^ or *Pi16*^+^) and pericytes (Pc; *Rgs5*^+^, *Abcc9*^+^), a rarer perineural cell type (PnC; *Wnt6*^+^, *Sfpr5*^+^) and other specialized fibroblasts^25–27^. ScRNA-seq analysis of lineage traced tdTom^+^ tongue cells, from *Hic1; tdTom* mice, identified these 3 cell types (Extended Data Fig. 2a and Supplementary Table 2). *Pdgfra*^+^ fibroblasts could be further sub-divided into 4 sub-types, Fb1-4: Fb1(*Pi16*^+^, *Cxcl13*^+^, *Ly6c1*^+^), Fb2 (*Col15a1*^+^, *Crabp1*^+^, *Rspo1*^+^, *Dkk3*^+^, *Lgr5*^+^), Fb3 (*Pi16*^+^, *Penk*^+^, *Apoe*^+^, *Gpx3*^+^) and Fb4 (*Col15a1*^+^, *Aqp1*^+^, *Rspo1*^+^, *Lgr5^+^*, *Dkk3*^+^). Spatial transcriptomic analysis with a 348-probe set (Supplementary Data 3) identified 3 Fb populations, Fb1-3. Fb1 and 3 mapped to the scRNA-seq dataset, whereas Fb2 appears to be a combination of Fb2 and Fb4, which have transcripts in common (Extended Data Fig. 2b-d and Supplementary Table 4). In comparison to Fb1 and 3, Fb2 displayed a distinct distribution subjacent to the epithelial basement membrane (Extended Data Fig. 2d). *Pdgfra* and *Rgs5* transcripts were found to be distributed throughout the tongue stroma (Extended Data Fig. 2d). Similar to the tibialis anterior SkM^26^, within the transverse and vertical muscle fascicles, tongue tdTom^+^ *Pdgfra*^+^ fibroblasts (nuclear GFP) were identified along with capillary-associated tdTom^+^ *Pdgfra*^-^ pericytes (Extended Data Fig. 2e). In the ventral lamina propria, numerous solitary tdTom^+^ cells were observed within the interstitium, subjacent to the basement membrane (Extended Data Fig. 2f). In the tongue, unlike other muscle compartments^26,27^, a large number, ∼30%, of tdTom^+^ cells were not perivascular (Extended Data Fig. 2g).

To refine the cell-of-origin within the *Hic1*^+^ compartment, scRNA-seq was used to profile end-point tongue tumours (Extended Data Fig. 3a). Three biological replicates (tSRC1-3) were profiled and they all mapped to a single tumour cell (Tc) cluster (Extended Data Fig. 3b). Further transcriptional profiling revealed that the tumours were *Pdgfra*^+^, *Rgs5*^-^, reflective of a fibroblast origin (Extended Data Fig. 3c). In the UMAP of tongue and tumour samples (tSRC1-3), the tumour cluster was partitioned into sub-groups, which could be partly distinguished based on the expression of: *Mki67* (proliferative population), *Nkd2*, *Epcam, Sox2* and *Sall3* (Extended Data Fig. 3d and Supplementary Table 5). *Axin2,* a transcript indicative of activated WNT/μ-catenin signaling, was expressed throughout the tumour cluster, consistent with a role for WNT signaling in SyS sarcomagenesis^33^. Single cell DNA sequencing of tSRCs revealed them to be polyclonal in nature^34^, and this is apparent in histological analyses, where well-defined and isolated tumours can be identified in the anterior region of the tongue and the ventral lamina propria (Extended Data Fig. 3e). Tumours are marked by an abundance of spindle shaped monophasic cells, a large area of poorly differentiated cells and small islands of biphasic tumour cells typically surrounded by PD cells (Extended Data Fig. 3e). For analysis of transcript distribution in histological samples, Tc-enriched genes (Supplementary Table 5) were used to generate 100 custom probes, which supplemented a Xenium pre-designed murine brain probe set (Supplementary Table 3). Spatial transcriptomic analysis of a tumour sample indicated the presence of multiple tumour clusters, along with non-transformed constituents, including normal epithelial (Ep), immune cells (IC), fibroblasts (Fb), endothelial and mural cells (En + MC) and peripheral nerve (PN) populations (Extended Data Fig. 3f-h and Supplementary Table 6). Notably, several interesting sub-populations emerged that can be mapped to specific histological features, in addition to aiding resolution of tumour sub-cluster identity and associated transcript features. The monophasic sub-type is defined by *Nkd2* expression, and can be further divided into *Nkd2*^+^ *Epcam*^-^ or *Epcam*^+^, and these could be further sub-divided based on *Sox2* expression (*Sox2^hi^*or *Sox2^lo^*) (Extended Data Fig. 3i). Conversely, poorly differentiated tumour cells are enriched for *Sall3* and *Pax2*, with reduced *Nkd2* expression (Extended Data Fig. 3e, h, I). Within these regions, small islands of *Epcam*^+^ *Sall3*^+^ pseudo-glandular structures (Extended Data Fig. 3e) are present, and are histologically similar to the human PD-Bi tumour sub-type.

ScRNA-seq analyses of stifle and tongue tumours revealed highly concordant biological replicates at similar anatomical sites, with tumour cells clustering based on location (Fig. 2a-d). Both stifle and tongue tumours exhibited a predominantly neural/developmental signature, with nuanced differences in gene expression. In comparison to stifle tumours, tongue tumours exhibited markedly higher expression of a subset of this signature, including *Alx3*, *En2*, *Foxe1, Pax6* and *Tbx1* (Fig. 2c and Supplementary Table 7). In contrast, stifle tumours contained a higher abundance of numerous *Hoxa*, *b*, *c* and *d* transcripts in comparison to tongue tumours (Fig. 2c and Supplementary Data 1 and 7). ScRNA-seq analysis of stifle tumours revealed two prominent clusters that could be partly distinguished by *Mgp* (Extended Data Fig. 4a-d). Consistent with tongue tumours, they shared similar expression profiles, with tumours being uniformly *Pdgfra*^+^, *Cd34***^-^**, and *Axin2*^+^, with variable expression of *Nkd2*, *Sall3* and *Epcam*, indicative of tumour subgroups (Extended Data Fig. 4a-d and Supplementary Table 8). Stifle tumours also contained an abundance of monophasic tumour cells with different morphological characteristics that could be visualized invading the local skeletal muscle (Extended Data Figure 4e). Moreover, spatial transcriptomic analyses of stifle tumours revealed a predominantly monophasic signature (Extended Data Figure 4f-i and Supplementary Table 9), with widespread expression of *Nkd2*, *Epcam* and *Sox2* (with small *Sox2* deficient areas) and limited punctate *Sall3* expression. To better define tumour phenotype-transcriptomic relationships, integrated analysis was performed across multiple spatial transcriptomic datasets (stifle n=5, tongue n=3) (Fig. 2e and Supplementary Table 10). Furthermore, the spatial analysis provided more granularity with respect to tumour heterogeneity. In this regard, a number of Mp subsets (with the exception of Mp2 and 3) that clustered in close proximity were identified. The unique properties of Mp2 and 3 could be robustly defined, in part, by expression of a limited number of transcripts, such as genes encoding select Semaphorins, central regulators of cell morphology and motility^35^ (Fig. 2f, g, i and Extended Data 4j). Three clusters exhibiting a PD signature (*Nkd2*^-^, *Sall3*^+^) were identified, and could be distinguished by *Hapln1* and *Troy* (*Tnfrsf19*) expression (Fig. 2g, i), a downstream target of the beta-catenin/WNT signaling pathway^36^. Moreover, transcriptomic signatures reflective of the different tumour sub-types could be detected to varying extents in tongue and stifle samples (Fig. 2h). Importantly, this integrated analysis reinforced the overarching transcriptomic features of the different tumour sub-types, and showed that *Nkd2* and *Sall3* identify distinct tumour cell domains, with *Sox2* exhibiting a gradient of expression from low (*Nkd2*^+^) to high (*Sall3*^+^) (Fig. 2j). It should also be noted that the *Sall3* and *Sox2* expressing subsets exhibited much higher *Mki67* expression compared to the *Nkd2* population. Collectively, these findings provide insights into the molecular basis of tumour heterogeneity and evolution, and suggest that this is related, at least in part, to the progressive acquisition of an earlier neural/developmental signature.

**Fig. 2:**
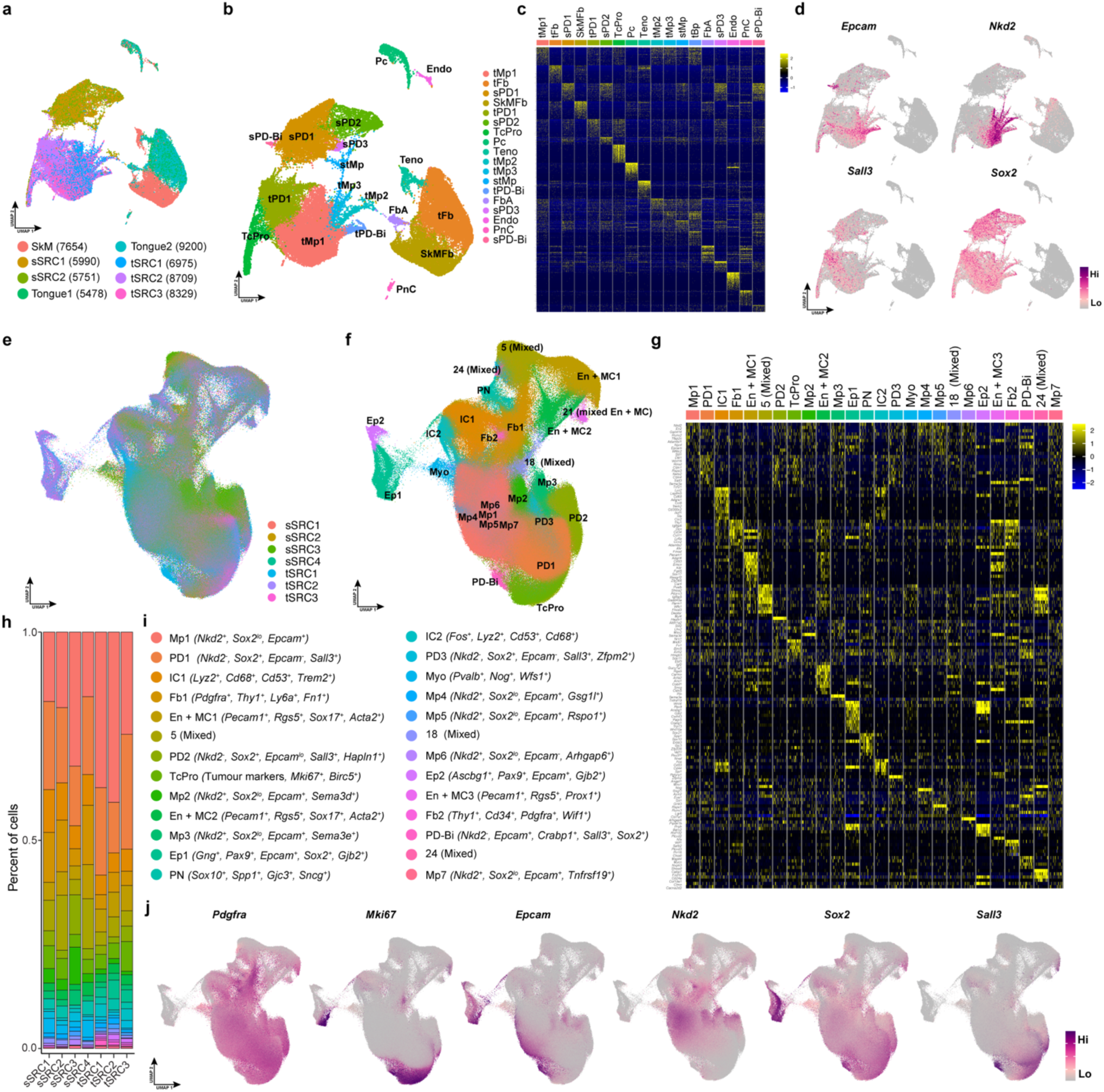
Molecular profiling of SyS tumours. **a,** UMAP of scRNA-seq data from the indicated samples, colored by sample ID. SkM, hind limb skeletal muscle with overlying facia; sSRC, stifle sarcoma; tSRC, tongue sarcoma. **b**, UMAP from (a), colored by cell cluster annotation with their assignment to different tumour phenotypes and cell types specified. t, tongue; s, stifle; SkM, hind limb skeletal muscle; Fb, fibroblast; Pc, pericyte; Teno, tenogenic; Endo, endothelial; PnC, perineural; Mp, monophasic; PD, poorly differentiated; PD-Bi, poorly differentiated biphasic; TcPro, proliferative tumour cell; FbA, activated fibroblast. **c**, Heatmap of scRNA-seq data showing enriched genes (top 50 upregulated transcripts) in the 18 clusters from (b). **d**, UMAPs colored by expression level of select transcripts. **e**, UMAP of integrated spatial transcriptomics analyses colored by sample ID (distinct biological replicates from samples in (a)). **f**, UMAP from (e) colored by cell cluster with cell types annotated based on markers listed in (i). **g**, Heatmap of the top 10 cluster marker gene expression per cluster. **h**, Stacked bar graph showing the percent distribution of clusters per tumour sample. **i**, Summary of markers that identify cluster cell types, including tumour subtypes. Cell type abbreviations as in c, with the addition of: IC, immune; En + MC, endothelial and mural; Ep, epithelial; PN, peripheral nerve; Myo, myofibre. **j**, UMAP from (e) colored by expression level of select transcripts.

## Kinetic analysis of SS18::SSX-induced reprogramming

The consistent latency and anatomical location of tongue tumours in the *Hic1; hSS2* model provided an opportunity to study the early stages of transformation. GFP^+^ cells were first detected using flow cytometry at 5 weeks post-induction with subsequent marked expansion (Fig. 3a and Extended Data 5a). In histological sections, small SS18::SSX^+^ lesions (< 30 cells) were identified at 4 weeks, indicating that immediately following *SS18::SSX* expression, there was a period of slow phenotypic modification prior to the emergence of a faster growing tumourigenic phenotype (Fig. 3a-b). These presumptive tumourigenic lesions were observed at the interface of the transverse and vertical muscle fascicles in the anterior tongue (Fig. 3b). As these tumours arise from the *Pdgfra*^+^ fibroblastic population, a *Pdgfra* allele containing H2B-EGFP (*Pdgfra^EGFP/+^*) was introduced into this background to improve visualization and capture of nascent tumour cells (Extended Data Fig. 5b). Haploinsufficiency of *Pdgfra* did not impact tumour latency nor phenotype, indicating that PDGFRA is not an SyS dependency (Extended Data Fig. 5c-f). Under these conditions, multiple individual tumour lesions were visualized starting at 5 weeks post-TAM, and likely coalesced into the larger tumour mass observed at end-point. To gain insights into these early stages, scRNA-seq analyses were performed on tongue *Pdgfra*^+^ cells from 5-days to 4-weeks after *SS18::SSX* induction and combined with later time-point datasets. Projection of these samples onto a force directed layout revealed the evolution of *Pdgfra*^+^ fibroblasts into a transformed phenotype (Fig. 3d-f). This was further explored in force directed layouts containing a subset of only *Pdgfra*^+^ cells (*Rgs5*^+^ and other minor contaminating cells removed) which revealed a linear trajectory from *Pdgfra*^+^ cells to tumour cells and loss of differentiation potential (Extended Data Fig. 6a-b). The initial stages of reprogramming were associated with fibroblast activation, and the expression of activation-associated transcripts such as *S100A4* increased, whereas quiescence-related transcripts, such as *Inmt*^26^, decreased (Extended Data Fig. 6c). Moreover, at the early stages, a separate, putative, non-productive trajectory emerged (Fig. 3g-h and Extended Data Fig. 6a), in which *Crabp1*, *Twist2*, and *Wif1* expression was elevated, along with other transcripts (Extended Data Fig. 6c). Shortly thereafter, numerous transcripts that define mature fibroblasts were down-regulated (i.e., *Cd34*, *Col15a1*, *Dpt*, *Inmt*, *Pi16*, *Ccl11*, *Ccl7*, *Thy1*) (Fig. 3e, h and Extended Data 6c). Productive transformation was further refined with pseudo-time analysis, in which numerous gene expression trends were identified. Transcripts exhibiting early expression in the transformation cascade included those encoding proteins involved in hedgehog and WNT signaling (Extended Data Fig. 6c). Subsequently, various transcripts involved in neurogenesis (i.e., *Gbx2*, *Nkx6-1*, *Pou3f1 and Pou3f2, Sox2 and Zic2*) and other developmental programs (i.e., *Barx2*, *Foxc2* and *Pax7*) emerged (Fig. 3e, h and Extended Data Fig. 6c). With time, transcripts reflective of more “advanced” tumours (i.e., *Epcam*, *Pax2, Mnx1, Olig1 and 2, Pou3f3 and Uncx*) were expressed (Fig. 3h, Extended Data Fig. 6c). Two tumorigenic trends were identified, distinguishing tSRC1 and 3 (*Sall3^lo^*) from tSRC 2 (*Sall3^h^*^i^) (Fig. 3g-h). Similar to that observed in the spatial transcriptomic analysis, *Nkd2^+^* populations appeared to be distinct from the *Sall3^+^* compartment, with both *Sox2^hi^* and *Sox2^lo^* groups (Fig. 3e). Moreover, *Epcam^+^* populations were also associated with *Nkd2^+^* and *Sall3^+^* compartments. Along this trajectory, numerous neurogenic factors were expressed, including Pou transcription factor encoding genes (i.e., *Pou3f1*-*3f3*), which play a prominent role in neurogenesis and the specification of neurogenic fates^37^. In this regard, SS18::SSX reprograms adult *Pdgfra*^+^ fibroblasts to a more developmental phenotype, with an abundance of active programs reflective of broad potential within the mesodermal and ectodermal lineages (Fig. 3i).

**Fig. 3:**
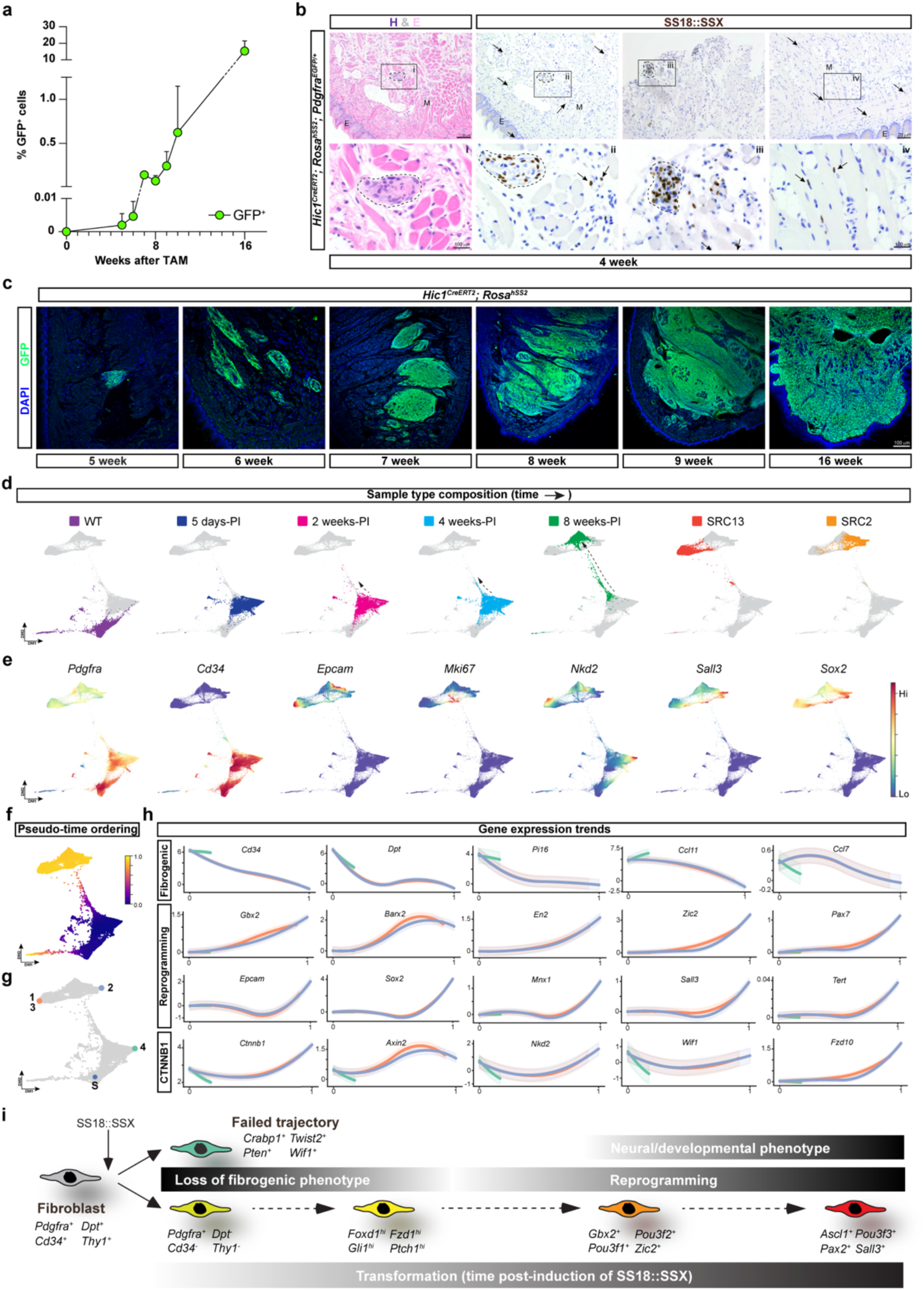
SS18::SSX-mediated reprogramming of *Hic1*^+^ *Pdgfra*^+^ MSCs. **a**, Quantification by flow cytometry of GFP^+^ cell abundance (expressed as percentage of total events) from tongue samples collected at various times post-TAM (n=3-4 per time point). **b**, Representative H&E images and corresponding anti-SS18::SSX staining (left two panels) on tongue tumours collected 4 weeks post-TAM administration. Additional anti-SS18::SSX stained samples are also shown (right panels). Dotted lines outline early SS18::SSX^+^ lesions. Arrows identify solitary SS18::SSX^+^ cells. M, skeletal muscle; E, epithelium. **c**, Representative epi-fluorescence images taken from tongue tumours collected at the indicated time points stained with anti-GFP. **d**, Force-directed layout (FDL) plots of scRNA-seq data from aggregated timepoints with individual time points highlighted by the indicated color within each panel. **e**, FDL plots from (d) colored by expression level of the indicated select genes. **f**, FDL plot from (d) colored by pseudo-time ordering of cells. **g**, Start cell and terminal end-point cells used for trajectory analysis and calculation of gene expression trends. S, start; 13, tSRCs 1 and 3 from Fig.2a; 2, tSRC2 from Fig.2a; 4, presumptive failed transformation protrusion. **h**, Gene expression trends of select transcripts derived from analysis of trajectories from (g). Trend line colours correspond to termini cells in (g) **i**, Summary schematic of SS18::SSX-induced fibroblast reprogramming including key inflection points.

## *Lgr5*^+^ fibroblasts and sarcomagenesis

*Hic1* is broadly expressed within the fibroblastic and pericytic compartments; however, in the tested genetic background, SyS tumours appeared at a high frequency at only 2 locations. Thus, it appeared that these two sites may harbour a *Hic1*^+^ *Pdgfra*^+^ population(s) and/or an environment that is more conducive to SS18::SSX-mediated transformation. To address the underlying basis for this selectivity, programs activated early in the reprogramming process were interrogated. Multiple components indicative of activated beta-catenin/WNT signaling (*Axin2*, *Ctnnb1*, *Fzd1, Fzd10*) appear early following *SS18::SSX* induction (Fig. 3h and Extended Data Fig. 6c). Numerous transcripts encoding factors involved in beta-catenin/WNT signaling were queried across tissues, and a prominent *Lgr5^+^* population was identified in the tongue, in comparison to other tissues (Extended Data Fig. 7a-b). *Lgr5* encodes a G-protein coupled receptor that acts to positively modulate beta-catenin/WNT signaling in the presence of its cognate ligands, the R-spondins^38^ (encoded by *Rspo* genes). *Lgr5* is expressed in numerous epithelial stem cell compartments, with limited expression in fibroblasts^39^. Previous studies described a rare *Lgr5*^+^ population within the tongue, particularly in the ventral lamina propria^40^ (Extended Data Fig. 7c), which overlaps with a *Hic1; tdTom*^+^ population (Extended Data Fig. 7d). Accordingly, *Lgr5* null mice present with ankyloglossia^41^. To better define this population, we introduced the *Pdgfra^EGFP/+^* allele into an *Lgr5^CreERT2/+^*; *Rosa^tdTom/+^* reporter background. Numerous lineage-traced *Lgr5*^+^ cells were identified in the intestinal epithelium; however, in this compartment, there was no overlap with *Pdgfra*^+^ cells (Extended Data Fig. 7e). In contrast, in the tongue, GFP^+^ tdTom^+^ cells were observed in the dorsal and ventral regions, subjacent to the epithelium (Extended Data Fig. 7f), consistent with that observed for the distribution of *Pdgfra* and *Lgr5* transcripts (Extended Data Fig. 7g). *Lgr5* and *6* are expressed with different kinetics following *SS18::SSX* induction, along with *Rspo3*, and *Lef1*, a major downstream effector of beta-catenin/WNT signaling^42^ (Fig. 4a). Tongue and stifle tumours exhibited uniform *Lgr5* and irregular *Lgr6* expression (Extended Data Fig. 8a-b). To address if this rare *Lgr5*^+^ population served as a potential cell of origin, *SS18::SSX* expression was induced in the *Lgr5^CreERT2^* background (Extended Data Fig. 8c). SyS tumours appeared in the tongue with a similar latency and phenotype to that observed with *Hic1^CreERT2^* (Fig. 1c, 4b-c). ScRNA-seq UMAPs showed that the transcriptomes of cells from the *Lgr5*-derived tumours co-clustered with *Hic1*-derived tSRC samples (Fig. 4d and Extended Fig. 8d), and expressed a similar constellation of hallmark transcripts (Fig. 4e and Extended Data Fig. 8e) and proteins (Extended Data Fig. 8f). Furthermore, human SySs were found to express *LGR5* and/or *LGR6,* and these two genes defined two stable epigenetic states^1^(Fig. 4f). Collectively, these findings indicate that at least a subset of SySs originate from an *Lgr5*^+^ *Pdgfra*^+^ fibroblastic population.

**Fig. 4:**
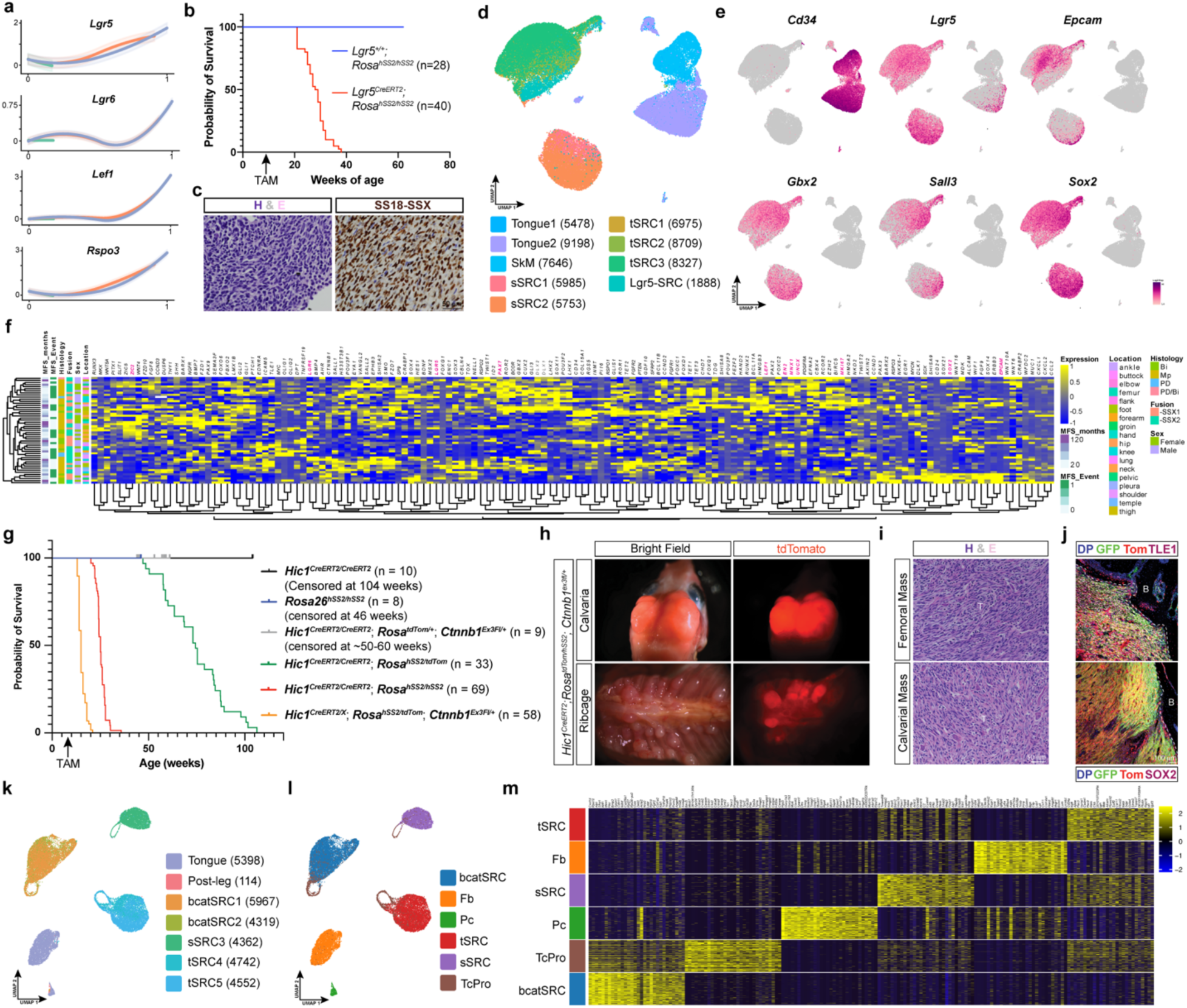
*Lgr5*^+^ MSCs and beta-catenin/WNT signaling in synovial sarcomagenesis. **a,** Gene expression trends of WNT signaling components (derived from pseudotime analysis in Fig. 3). **b**, Kaplan-Meier survival curve of indicated genotypes, with TAM administration at ∼8 weeks of age. **c**, Representative images of H&E and anti-SS18::SSX stained sections of tongue tumours derived from *Lgr5^CreERT2^; Rosa^hSS2/hSS2^* mice. **d**, UMAP colored by the indicated scRNA-seq sample IDs. **e**, UMAP from (d) colored by expression level of the indicated select transcripts. **f**, Heatmap of differentially expressed genes from human SyS samples. Differentially expressed genes were identified using the mouse dataset and expression pattern queried across the human data. Metadata is colour coded and includes metastatic free survival in months (MFS_months), MFS_event, histological sub-type (Bi, biphasic; Mp, monophasic; PD, poorly differentiated), nature of the fusion (*SS18::SSX1* vs *SS18::SSX2*), sex and tumour location. **g**, Kaplan-Meier survival plots of the indicated genotypes. **h-j**, Representative images of peri-skeletal tumours in the specified genetic background. **h,** whole-mount images of tumours from the indicated sites. **i**, H&E-stained tumour sections from the indicated sites. **j**, Immunofluorescence images of anti-TLE1 and anti-SOX2 stained tumours from the indicated sites. **k**, UMAP of scRNA-seq colored by sample ID. **l**, UMAP from (k) colored by cluster identity. **m**, Heatmap of the top 10 cluster markers for the clusters identified in l.

LGR5 acts to promote beta-catenin/WNT signaling^38^, suggesting that the transformation supportive nature of the *Lgr5*^+^ compartment may be related to higher basal beta-catenin/WNT signaling. Consistent with this premise, previous studies have demonstrated a dependency for beta-catenin/WNT signaling in synovial sarcomagenesis^33^, and SyS formation was substantially increased in the presence of an activating allele of beta-catenin^43^ (*Ctnnb1^Λ1ex3^*). *Ctnnb1* is also a dependency in human SyS cell lines^44^ (Extended Data Fig. 8g). To further evaluate this, we introduced the *Ctnnb1^ex3floxed/+^ allele* into the *Hic1^CreERT2^*; *Rosa^hSS2/tdTom^* background (Extended Data Fig.8h). In the absence of TAM, no notable phenotypes were observed; however, after TAM administration, mice in the *Hic1^CreERT2^*; *Rosa^hSS2/tdTom^*; *Ctnnb1^ex3floxed/+^* group reached clinical end-point with a median latency of ∼7 weeks post-induction (Fig. 4g). Necropsy revealed a high tumour burden, with numerous TLE1 positive tumours, characteristic of monophasic SyS, decorating the skeletal system (Fig. 4h, i, j). Transcriptomic scRNA-seq profiling revealed that tumours in this background clustered separately from stifle and tongue tumours (Fig. 4k-m). *Ctnnb1^Λ1ex3/+^*-derived tumours abundantly express *Pdgfra*, *Lgr5* and other beta-catenin/WNT signaling components (Extended Data Figure 8i-l), as well as, markers reflective of the earlier stages of reprogramming (i.e., *Gbx2*), but lacked transcripts indicative of intermediate or “more reprogrammed” cells (i.e., *Epcam*, *Mnx1*, *Sall3*, *Sox2*) (Fig. 4m, Extended Data Fig. 8l and Supplementary Table 11). However, these tumours retained competency to express various cytokines as exemplified by *Cxcl5* expression (Extended Data Fig. 8l). Collectively, these findings suggest that canonical WNT/beta-catenin signaling contributes to a fibroblast state amenable to SS18::SSX transformation; however, high constitutively active beta-catenin/WNT signaling likely interferes with complete fibroblast reprogramming.

## SS18::SSX-induced epigenomic reprogramming

Stifle and tongue tumours exhibit concordant and consistent transformation trajectories, resulting in a unique neural/developmental transcriptome that is a hallmark of human SyS^22^. SS18::SSX has been shown to modify the epigenome and interact directly with the histone modification H2AK119ub, a mark deposited by the PRC1.1 complex that is typically associated with transcriptional repression^10,15–17^. To gain additional insights into the underlying basis of the SyS neural/developmental program, a series of histone modifications associated with gene activation and repression, along with chromatin accessibility, were assessed in tongue derived *Hic1; tdTom* WT MSCs and *Hic1; hSS2* SyS tumour cells. ScATAC-seq revealed that even at 8 weeks post-induction, sarcomagenesis activated transcripts (SAT) exhibited open chromatin domains, whereas sarcomagenesis silenced transcripts (SST) exhibited limited accessibility (Fig. 5a and Extended Data Fig. 9a). It became clear that SS18::SSX modifies the *Hic1*^+^ fibroblast epigenome in a deliberate pattern, but why these specific loci were targeted, remained unanswered. ChIP-seq profiling revealed several interesting features (Fig. 5b and Extended Data Fig. 9b), including a marked decrease of H3K27me3 observed at promoter regions in SyS tumours compared to WT MSCs (Fig. 5b). To examine this further, histone modifications at SAT, maintenance transcripts (MAT, i.e., expressed but not changed following transformation) and SST genes (Supplementary Table 12) were analyzed, and showed that SAT genes in WT cells were associated with a bivalent signature (H3K4me3 and H3K27me3), whereas MAT genes revealed no notable pattern differences (Fig. 5c-d). In contrast, at *Cd34* (SST), transformed cells exhibited a loss of H3K4me3 and a gain of H3K27me3 compared to WT (Fig. 5d). These loci also showed differential occupancy of H2AK119ub, with the SAT genes demonstrating pronounced enrichment of this mark in comparison to MAT and SST loci (Fig. 5c, d), and this was coupled with selective occupancy of SS18::SSX in these regions^45^. SAT genes include many of the *Hox* genes in clusters *A*, *B*, *C* and *D*, and these loci also exhibited similar patterns of bivalency with H2AK119ub (Extended Data Fig. 10a). Analysis of histone marks at single loci (*Hox* clusters, *Sall3*, *Sox2* and *Cd34*) in WT cells showed that *Hox* genes, *Sall3* and *Sox2* (SATs) exhibit similar H2AK119ub, H3K4me3, H3K27me3 peaks around the promoter, which extend into the gene body (Fig. 5d and Extended Data Fig. 10b). At these loci, following transformation, H3K27me3 peaks declined and the H2A119ub peak was diminished, albeit still detectable. SST loci were typically associated with either weak or absent H2AK119ub marks in either WT or transformed cells (Fig. 5c). Collectively, these findings indicate that SAT genes exhibit a specific pattern of bivalency, along with H2AK119ub, in the cells of origin, and that these marks resolve to H3K4me3 and increased transcription following SS18::SSX-mediated epigenomic reprogramming. This pattern of histone modification is further confirmed in the analysis of 52 human SySs, which showed a continuum of bivalency at overlapping loci to murine SySs, with preferential loss of H3K27me3 and resolution to predominantly H3K4me3^1^.

**Fig. 5:**
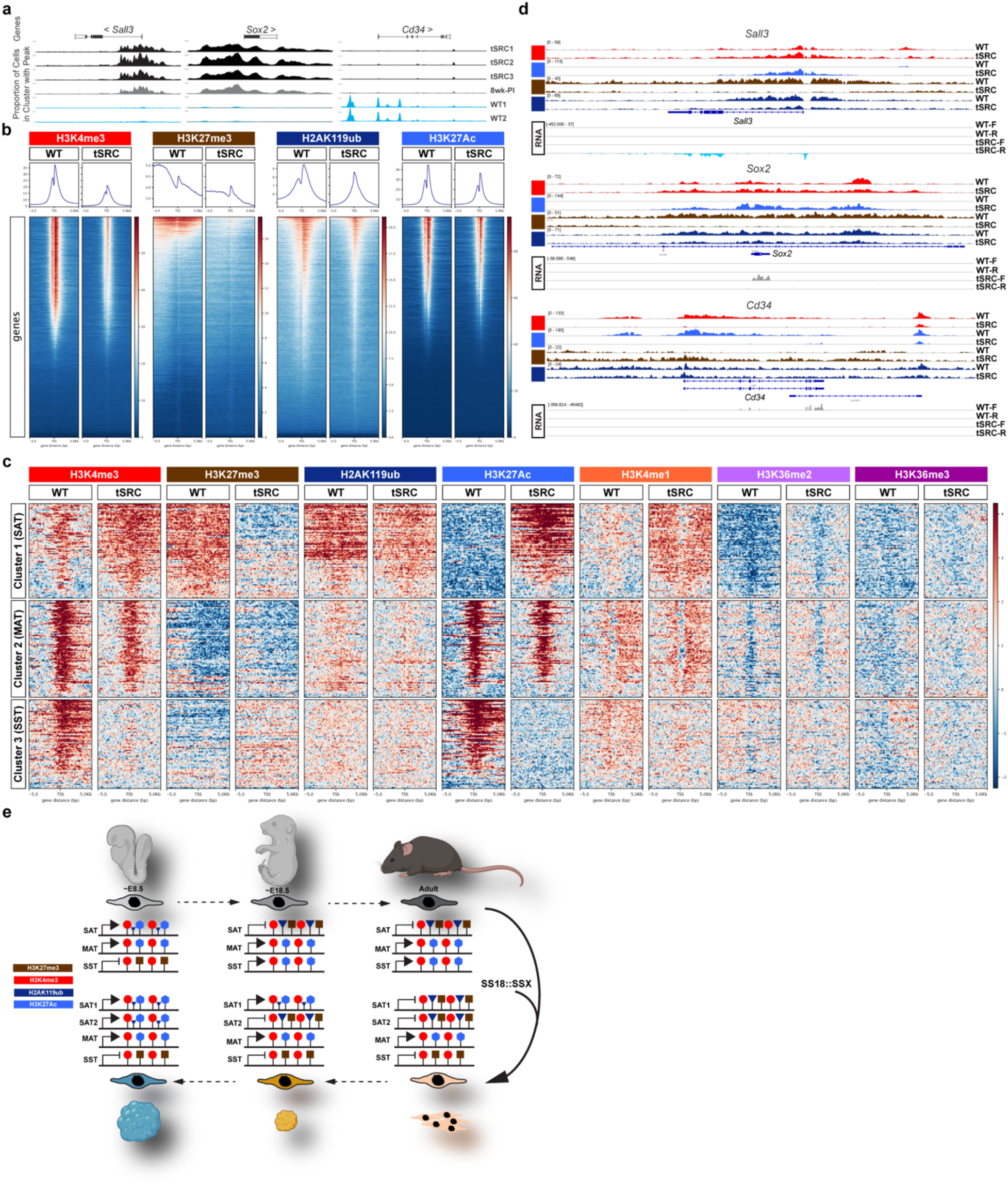
Epigenetic profiling reveals a H2AK119ub and bivalent signature underlying SS18::SSX-mediated transformation. **a,** Gene track plots derived from scATAC-seq analysis of the indicated tongue samples. **b**, Profile heatmaps of the indicated histone modifications around the TSS of RefSeq genes. The gradient, blue-to-red, represents low-to-high counts in the noted regions. **c**, Heatmaps of indicated histone modifications for select sarcomagenesis activated (SAT), maintenance (MAT) and sarcomagenesis silenced (SST) transcripts, from 75 loci (TSS +/- 5kB) each. **d**, Gene track plots of ChIP-seq peaks for the indicated histone modifications and RNA-seq reads, detected at the indicated loci. Tracks are colored according to marks in (b) and (c). WT, wildtype tongue MSCs; tSRC, transformed cells from endpoint tongue tumors; F and R denote reads from the + and – strand, respectively. **e**, Model of SS18::SSX-mediated transformation of fibroblasts and their acquisition of a more embryonic phenotype.

The neural/developmental signature of SyS appeared to be embedded in the epigenome of adult *Hic1*^+^ MSCs and to explore the origin of this program, embryonic MSCs (embryonic age E11.5 – E16.5) were profiled by scRNA-seq (E11.5 – E16.5) and ChIP-seq (E12.5) (Extended Data Fig. 11a). Through development, MSCs acquire a more “adult-like” phenotype, which is associated with up-regulation of numerous genes at ∼E16.5 that are reflective of mature fibroblasts (*Ccl2*, *Ccl7*, *Ccl11*, *Cd34, Col15a1*, *Dpt*, *Cd34, Il6, Inmt, Pi16*), and down-regulation of genes linked to an embryonic multipotent MSC phenotype (i.e., *Id* genes, *Hmga2, Sall3, Shisa2*, *Sox11*) (Extended Data Fig. 11b and d). In contrast, transformation of adult WT to tSRC cells exhibited an anti-parallel trajectory, whereby mature fibroblast markers are down-regulated and transcripts reflective of embryonic MSCs are up-regulated (Extended Data Fig. 11b). Similar to that observed at SAT loci in adult WT MSCs, embryonic MSCs (E12.5) exhibited bivalency and H2AK119ub at these loci (Extended Data Fig. 11c-d). This pattern is not present at SST genes (Extended Data Fig. 11 c-d). Taken together, the neural/developmental signature of SyS, is reflective of the embryonic heritage of the cell of origin, which is efficiently unmasked following SS18::SSX-mediated reprogramming.

## Discussion

Sarcomagenesis provides a unique opportunity to study the mechanisms underlying cell transformation, as these tumours are typically genetically quiet and have single, well-defined drivers, as is the case with SS18::SSX and synovial sarcoma^1^. Many sarcomas are presumed to arise from tissue-resident mesenchymal cells, and here we show that expression of *SS18::SSX* in *Hic1*^+^ fibroblasts leads to robust SyS generation, suggesting that these cells represent strong origination potential for SyS. The ability to follow tumorigenesis in this model from inception of oncoprotein expression through to tumours has enabled us to further refine the cell of origin and delve into the SS18::SSX-induced reprogramming cascade. In this manner, we show that SyS tumours arise from a rare *Hic1*^+^ *Pdgfra*^+^, *Lgr5*^+^ fibroblast subset, and that these tumours also widely express *Lgr6*. Trajectory and expression analysis indicates that *Lgr6* expression is subsequent to *Lgr5*, and thus, *Lgr5* and *Lgr6* may define distinct stages along a transformation continuum. Alternatively, *Lgr6*^+^ fibroblasts may serve as an additional source of SS18::SSX-transformation competent fibroblasts. Consistent with this, unsupervised analysis of human tumour transcriptomic and epigenomic data led to the identification of two major sub-groups, distinguished by *LGR5* and/or *LGR6* expression^1^.

SS18::SSX reprogramming of *Hic1*^+^ fibroblasts is comparable to that observed with the generation of induced pluripotent stem cells (iPSCs) following expression of *Klf4, Myc, Oct4 (Pou5f1)* and *Sox2* (OSKM)^46^. The early stages of both programs are associated with a loss of a fibrogenic phenotype and the subsequent progressive acquisition of a transcriptome and epigenome reflective of earlier stages of embryogenesis, marked by widespread loss of H3K27me3^47–50^. With the exception of *Pou5f1*, SySs often express the 3 remaining factors, which have proved insufficient for complete reprogramming of fibroblasts to iPSCs^46^. During OSKM-mediated reprogramming, several groups^51^ identified a partially reprogrammed hyperproliferative population that was associated with the expression of a set of transcripts, including *Sall3*, *Ascl1*, *Neurog2*, *Neurog3*, *Isl1*, *Prox1*, *En2* and *Foxd1*^49^. With the exception of *Isl1* which is not expressed in murine SyS, but does exhibit promoter bivalency, the rest of these transcripts represent SATs. Numerous transcripts associated with pluripotency (i.e., *Pou5f1*, *Nanog*) are not expressed in SyS, nor do their promoters exhibit bivalent marks, indicating that the extent of SS18::SSX-mediated reprogramming is likely hardwired into the fibroblast precursor epigenome. Collectively, these findings suggest that SS18::SSX drives tumorigenesis through the forced induction and maintenance of an epigenome-embedded embryonic transcriptional program (Fig. 5e).

Following SS18::SSX induction, there appears to be a period of slow limited growth (∼4-5 weeks), after which tumour cell proliferation increased substantially, such that ∼21% of the cells in end-point tumors were cycling. This slow period likely involving chromatin remodelling, is also congruent with that observed in iPSC generation from fibroblasts^52^. During this period numerous fibroblast-expressed cytokines continue to be expressed in SS18::SSX-expressing fibroblasts, and shortly after SS18::SSX induction, chemokines associated with fibroblasts activation^26^ are also transiently increased (i.e., *Cxcl5*). However, with continued reprogramming to an “earlier” embryonic state, competency to express a large number of pro- and anti-inflammatory cytokines (i.e., *Ccl2*, *Ccl11*, *Il6*, etc.) is lost. In the developing murine embryo, cytokine expression in fibroblasts starts to become detectable at ∼E16.5^53^. Thus, the apparent uptick in tumor growth, is coincident with the expression of more potent growth-promoting embryo-associated transcription factors. However, a contributing factor may also involve the widespread loss of cytokine transcriptional competency and the emergence of a cold immune phenotype, reflective of human tumours^54,55^. With regards to the former, SyS tumours express a large repertoire of “embryonic” transcription factors, any one of which alone or in combination, is sufficient to promote a tumourigenic phenotype. Differential expression of these factors is linked to tumor heterogeneity and the extent of reprogramming. Moving forward, the *Hic1; hSS2* mouse model will be a helpful tool to study how this progressive process unfolds.

## Acknowledgements

This work was supported by the following grants: National Institutes of Health grants CA231652 (K.B.J., M.H., T.O.N., T.M.U.), Terry Fox New Frontiers Program Project Grant (1021, M.H., T.O.N., T.M.U.) and Canadian Institutes of Health Research (PT-025845, M.H., T.O.N., T.M.U.). M.A. was supported by a UBC graduate scholarship. We would like to thank Dr. Timothy O’Connor for helpful discussions. We would also like to thank the following facilities, UBC Epigenetics Core, BRC-seq, ubcFLOW cytometry, AbLab, BRC genotyping, Ingrid Barta of the ACS Diagnostic and Research Histology Laboratory Services and the BRC transgenic unit.

## Lead contact statement

Correspondence and requests for materials should be addressed to T. Michael Underhill.

## Author contributions statement

L.A.H., R.W.S., L.A.M., M.A. and G.D. were responsible for performing all experiments. L.A.H., R.W.S., M.A., M.V. and J.H. were involved in bioinformatic analyses. X.Q.W. and T.O.N. were responsible for histological scoring. L.A.H., R.W.S., L.A.M., M.A., J.L., K.B.J., M.H., T.O.N. and T.M.U. participated in experimental design, data interpretation and preparation of the manuscript. All authors provided feedback and editing of the manuscript.

## METHODS

### Mice

Mice were housed under standard conditions (12 hr light/dark cycle) and provided food and water *ad libitum*. For all experiments, litter mates of the same sex were randomly assigned to experimental groups. Animals were maintained and experimental protocols were conducted in accordance with approved and ethical treatment standards of the Animal Care Committee at the University of British Columbia.

Generation of the *Hic1^CreERT2^* mouse line (Jax 036704) has been previously described^34^, and briefly, involved the insertion of an IRES-CreERT2 cassette into the 3’ UTR of the *Hic1* gene. For all experiments, *Hic1^CreERT2/CreERT2^* mice were used and will be referred to as *Hic1^CreERT2^*. Other mouse lines include: B6.Cg-Gt(ROSA)26Sor^tm14(CAG-tdTomato)Hze^/J (herein referred to as *Rosa^td^*^Tom*/+*^; Jax stock number 007914), C57BL/6J (Jax stock number 000664), *B6.129P2-Lgr5^tm1(cre/ERT2)Cle^/*J (*Lgr5^CreERT2^*; Jax 008875), B6.129S4-*Pdgfra^tm11(EGFP)Sor^*/J (*Pdgfra^EGFP^*; Jax 007669), *B6.Cg-Ndor1^Tg(UBC-cre/ERT2)1Ejb^/1J* (UBC-CreERT2, Jax 007001), *Ctnnb1^lox(e3)^* (*Ctnnb1^delta3floxed^*)^56^ and *Rosa^hSS2^* (conditional allele of *SS18::SSX2* with a downstream IRES-EGFP cassette inserted in the *Rosa26* locus)^30^. With the exception of the *Ctnnb1^delta3floxed^* line, all animals used in these experiments were in a C57BL/6 background.

For tumour studies, *Hic1^CreERT2^*, UBC-CreERT2 or *Lgr5^CreERT2^* were interbred with *Rosa^hSS2^* to generate mice either heterozygous or homozygous at the *Rosa^hSS2^* locus. *Hic1^CreERT2^; Rosa^hSS2/hSS2^* mice will be referred to as *Hic1; hSS2*, and their non-tumorigenic control counterpart, *Hic1^CreERT2^; Rosa^td^*^Tom*/+*^ are indicated as *Hic1*; *tdTom*. To manipulate the canonical WNT pathway during concomitant *SS18::*SSX2 expression, *Hic1^CreERT2^*; *Rosa^hSS2/tdTom^* were interbred with the *Ctnnb1^delta3floxed/+^* line to generate *Hic1^CreERT2^*; *Rosa^hSS2/tdTom^*; *Ctnnb1^delta3floxed/+^* mice. In this background, the presence of the conditional tdTomato (tdTom) allele enabled lineage tracing. To efficiently identify Pdgfra^+^ fibroblasts, a single allele of *Pdgfra^EGFP^* was introduced into the *Hic1^CreERT2^*; *Rosa^hSS2/hSS2^* background.

For genotyping, ear punches were obtained at 3-8 weeks of age. DNA extraction and genotyping were performed using polymerase chain reactions by the Biomedical Research Centre Genotyping Facility, using the primers listed in Supplementary Information. To induce CreERT2 nuclear translocation, 8–9-week-old mice were intraperitoneally (IP) injected with 0.1 mL of 25 mg/mL tamoxifen (TAM; Sigma, T5648) in corn oil (Sigma, C8267), every 24 h for 5 consecutive days.

### Immunofluorescence, immunohistochemistry and histology

To enable detection of native EGFP and/or tdTomato expression in processed tissue samples, mice were terminally anesthetized by IP injection of Avertin (400 mg/kg; Sigma, T48402), and fixed by intracardial perfusion of 10 mM EDTA in PBS, followed by 2% paraformaldehyde (10-15mL each). Tissues were immersed in 2% paraformaldehyde for 48-72 h at 4°C, in the dark.

For cryosectioning, fixed samples were washed 3 x 30 min with PBS, and then incubated through a cryoprotective series of sucrose solutions of increasing concentration (10-50%), at 4°C for ≥ 3 h each. Tissues were embedded into disposable plastic cryomolds (Polysciences, 18646A) containing OCT (Tissue Tek 4583), and frozen in an isopentane bath cooled by liquid nitrogen. Cryosections were cut (Leica CM3050S or CM1950) at a thickness of 5-30 μm and mounted onto Superfrost Plus slides (VWR, 48311-703).

For immunofluorescence (IF) staining, slides were thawed at room temperature, washed 3 x 10 min in PBS and incubated for 1 h in PBS containing 10 mg/mL sodium borohydride (Sigma, 213462) to quench autofluorescence. Slides were then rinsed 2 x 1 min with PBS, and incubated in block solution containing 2.5% BSA (Sigma, A7030) and 2.5% goat serum (Gemini, 100-190) for 90 min at room temperature, prior to incubation in primary antibody overnight at 4°C. The following primary antibodies were used: α-CD31 (1:50, 550274, BD Biosciences), α-RFP (1:100, ab62341, Abcam), α-GFP (1:100, ab13970, Abcam), a-SOX2 (1:100, ab97959, Abcam), α-OLIG2 (1:100, ab9610, Sigma), α-Laminin (1:100, Ab11575, Abcam), α-TLE1 (1:50, Ab183742, Abcam), α-MUC1 (1:50, Ab15481, Abcam), α-SS18::SSX (1:100, 72364S, Cell Signaling). EGFP fluorescence signal from the hSS2 allele was detected using anti-GFP for IF image panels. Controls consisted of incubation in the absence of primary antibody. Alexa Fluor conjugated secondary antibodies (ThermoFisher) were diluted 1:500 and applied to slides for 45 min at room temperature in the dark. Secondary antibodies included: Alexa Fluor 488 (A11039), Alexa Fluor 647 (A21247), Alexa Fluor 647 (A21245), Alexa Fluor 555 (A21434) and Alexa Fluor 594 (A11012). After each antibody incubation, 3 x 5 min PBS washes were performed. Following application of secondary antibody, sections were counterstained with DAPI (600 nM in PBS for 5 min), rinsed 2 x 2 min with PBS and mounted with Aqua Polymount (Polysciences, 18606).

For immunohistochemistry (IHC) and H&E analyses, following dissection, tissues were fixed in 10% neutral buffered formalin for 24-72 h, washed 3 x 30 min in PBS, then placed in 70% ethanol. Paraffin embedding, sectioning, and H&E staining were performed according to standard protocols at the UBC ACS Diagnostic & Research Histology Laboratory. FFPE tissues were also utilized for SS18::SSX IHC, as previously described^57^. Briefly, 4 um sections were subjected to heat-induced antigen retrieval for 20 min, followed by SS18::SSX antibody (1:300, 72364, Cell Signalling Technology) incubation for 15 min at 37°C. Visualization was achieved using the BOND Polymer Refine Detection kit (DS9800, Leica Biosystems).

### Image acquisition and quantification

Bright-field and wide-field epi-fluorescence imaging was performed on an Olympus BX63 microscope with an Olympus DP72 or XM10 camera, and images were acquired using the Olympus CellSens Program. In addition, bright field images were also acquired using the ZEISS Axio Scan and Aperio ImageScope software (Leica Biosystems). Confocal images were acquired using either a Nikon Eclipse Ti inverted microscope equipped with a C2Si confocal system or Nikon Ti-E inverted microscope with an A1R HD25 confocal scanning head at 1024 x 1024 resolution, with 4x average. Maximum intensity projections were subsequently generated and denoised using the *denoise.ai* function from the GA3 module of the NIS elements image analysis software (Version AR 5.41.02). Monochrome TIF images were imported into Fiji (ImageJ2; version 2.9.0/1.53t) to merge channels and adjust the brightness and contrast (LUTs) of images to comparable levels.

To evaluate Hic1^+^ MSC distribution with regards to vessel proximity, tongues from *Hic1; tdTom* control mice (n=3) were collected and processed for IF with CD31 antibody staining, as described. Three 20X images were taken from separate, non-adjacent sections from each animal. All tdTom^+^ cells were counted in each image and evaluated for their vessel association status. Cells were deemed vessel-associated when they were contacting a CD31-positive vessel and deemed non-vessel-associated when they had no clear contact. The mean of all image counts was calculated and vessel-association was expressed as a percentage of the total number of tdTom^+^ cells (DAPI^+^).

### Tumour type counting

Animals were sacrificed at humane endpoint and necropsies performed. Lesions were identified by gross examination and validated using GFP fluorescence on an SMZ25 stereo dissection microscope (Nikon). Anatomical locations of GFP^+^ lesions were recorded and tabulated.

### Cell cycle analysis

To enable detection of proliferating cells, standard methodology involving incorporation of the nucleoside triphosphate analogue 5-ethynyl-2’-deoxyuridine (EdU) was used. For these purposes, animals were injected IP with 0.5 mg of EdU (ThermoFisher, E10415) in 250 uL PBS, daily, for 5 days. All experimental (*Hic1; hSS2*, n=6) and control (*Hic1; tdTom*, n=3) animals were injected with EdU beginning at 23 weeks of age (14 weeks post-TAM treatment). Histological samples were collected 24 h after the last EdU injection, and EdU^+^ cells were detected using the Click-iT Plus EdU Alexa Fluor 647 Imaging kit (ThermoFisher, C10640), as per manufacturer instructions. After completion of the Click-iT reaction, the slides were stained with the desired immunofluorescent antibodies (as described, starting at the sodium borohydride step). Three 20X images were taken from separate, non-adjacent sections in each animal. For control animals (n = 3), all tdTom^+^ and tdTom^+^EdU^+^ cells were counted in each image, the mean of all image counts was calculated, and the proliferation rate was expressed as a percentage of the total number of tdTom^+^ cells. For experimental animals (n = 6), all GFP^+^ and GFP^+^EdU^+^ cells were counted in each image, the mean of all image counts was calculated, and the proliferation rate was expressed as a percentage of the total number of GFP^+^ cells.

### Population RNA-sequencing of tumours

Tumours and control tissues (comparable anatomical region) were harvested and dissociated using a hand-held tissue homogenizer (Omni International Inc.) in 2-3 ml RNAzol (Sigma, 4533). Total RNA was isolated using RNAzol, as per the manufacturer’s instructions. RNA quality was assessed with an Agilent Bioanalyzer 2100 RNA 6000 Nano kit. Libraries were generated from RNA samples with an RNA Integrity Number > 8, using the standard TruSeq Stranded mRNA library kit protocol. Paired end sequencing was performed on the Illumina NextSeq 2000 using the P3 300 cycle kit.

### RNA-seq Bioinformatic Analyses

Illumina sequencing outputs generated bcl files that were demultiplexed by bcl2fastq2. Demultiplexed read sequences were aligned to the Mouse Genome (mm10) reference sequence using TopHat^58^ splice junction mapper with Bowtie 2^59^ or STAR^60^ aligners (RNA-seq Alignment app, Illumina Basespace). Assembly was performed and differential expression were estimated using Cufflinks^61^ and Cuffdiff^62^, respectively (Cufflinks Assembly & DE app; Illumina Basespace). For robust identification and statistical validation of differentially expressed genes, gene sets were also analyzed for differential expression using edgeR, DESeq2 and manually cross referenced with single cell data and differential expression results from the *cellranger* pipeline analysis. Hierarchical clustering of population RNA-seq data and heatmap generation were performed with the pheatmap R package using Euclidian distance.

### Flow cytometry and fluorescence-activated cell sorting (FACS) methodology

Tissues from *Hic1; tdTom* control mice (tongue), *Hic1; hSS2* experimental mice (tongue and stifle masses) with and without *Pdgfra*^EGFP/+^, *Hic1^CreERT2^; Rosa^tdTom/hSS2^; Ctnnb1^deta3floxed/+^*experimental mice (skeletal masses) and *Lgr5^CreERT2^*; *Rosa^hSS2/hSS2^*experimental mice (tongue and lip masses) were processed using the same protocol. After dissection, tissues were minced with scissors and placed in Collagenase II (500u/mL, Sigma, 6885) digest cocktail (activated with 10μL/mL 250mM CaCl2) for 30 min at 37°C. Tissues were washed twice in ice-cold PBS, with centrifugation at 120g for 1 min at 4°C. After the second centrifugation, tissue pellets were resuspended in Collagenase D (1.5u/mL, Roche, 11088882001)/Dispase II (2.4u/ mL, Roche, 04942078001) digest cocktail (activated with 10μL/mL 250mM CaCl2) and rotated for 1 h at 37°C with gentle vortexing every 15 min. After enzymatic digestion, the cells were washed in fluorescence-activated cell sorting (FACS) buffer (2% Fetal Bovine Serum in PBS with 2mM EDTA), filtered through a 40 μm cell strainer (Fisherbrand, 22363547), and centrifuged at 500g for 5 min at 4°C. The supernatant was discarded, and the cell pellets were resuspended in antibody cocktail (in FACS buffer) and incubated in the dark for 30 min on ice. To enrich for tdTom^+^ and/or GFP^+^ cells, a FACS strategy was employed with the following lineage (Lin) antibody cocktail: anti-Ter119-647 (1:200, 67-0031-01, Ablab), anti-CD31-APC (1:400, BD Biosciences, 551262), anti-CD117-APC (1:500, eBioscience, 17-1172-82), anti-CD11b-647 (1:500, Ablab, 67-0055-01), anti-F4/80-647 (1:500, Ablab, 67-0035-05) and anti-CD45-647 (1:400, Ablab, 67-0047-01). After antibody staining, the cells were washed in FACS buffer, centrifuged at 500g for 5 min at 4°C, resuspended in FACS buffer containing 4 μM Hoechst 33342 (Sigma, B2261) and 1.5 μM Propidium Iodide (Sigma, P4864), and transferred to 5 mL polypropylene round-bottom tubes with a cell-strainer cap (Falcon, 352063).

Samples prepared for FACS were enriched using a BD Influx cell sorter, with a combination of Hoechst, propidium iodide and forward/side scatter parameters used to identify viable cells. Control tongues and calvarial masses from the stabilized β-catenin model were sorted for tdTom^+^/Lin^-^ cells. Tongue and stifle tumours collected 8 weeks post TAM, and beyond, were sorted for GFP^+^/ Lin^-^ cells. As GFP expression from the hSS2 allele was relatively weak at earlier timepoints, samples collected prior to 8 weeks were from *Hic1*; *hSS2* animals containing the *Pdgfra^EGFP/+^* allele, and cells were sorted for GFP^+^/ Lin^-^ cells. Lgr5^CreERT2^ driven sarcomas were also sorted for GFP^+^/Lin^-^ cells, with the GFP signal originating from hSS2-EGFP and Lgr5-EGFP, or both.

All samples were sorted a second time using the same parameters to increase purity and viability. Sorted cells were collected into sort media (20% FBS, L-glutamine (2 mM), penicillin (100 units/ml), and streptomycin (100 μg/ml) in DMEM) in cooled siliconized microcentrifuge tubes (Fisher Scientific, 02-681-320). Cells were sorted with gating applied using BD FACS Software. For sequencing applications, sorted cells were visually evaluated and manually counted using a hemocytometer, prior to use for sequencing analyses (i.e., scRNA-seq, ChIP-seq, etc.).

For time course analyses of GFP+ cells, *Hic1*; *hSS2* animals were utilized and a mononuclear cell suspension was isolated from the tongues, as described above, at 5, 6, 7, 8, 9, 10 and 16 (endpoint) weeks (n=3-4 per timepoint). Cells were analyzed using a CytoFLEX flow cytometer (Beckman Coulter). First, cells were gated for mononucleated (Hoescht^mid^) Lin-cells, and then the percentage of GFP+ cells were enumerated. Fluorescence minus one controls were used for all experiments. FlowJo was used to analyze and visualize all cytometer and FACS data.

### Single cell RNA-Seq

Single cell suspensions were generated and tdTom^+^ and/or GFP^+^ cells were enriched by FACS (as described above) into DMEM containing 5% FBS. Cells were counted and quality control was evaluated by hemocytometer. If the single cell suspension was confirmed to contain >98% tdTom^+^ and/or GFP^+^ cells, the solution was input into a Chromium Controller (10X Genomics). Cell capture and library generation was carried out with the Chromium single cell 3’ reagent kit v2 or v3 (10X Genomics). cDNA libraries were sequenced on a Nextseq500 or 2000 (Illumina) to a depth > 40,000 reads per cell. The generated reads were aligned to a modified mm10 reference genome that contains the sequences for tdTomato and GFP. Demultiplexing and sequence alignment was performed using the cellranger *count* pipeline (10X Genomics). Aggregated libraries were generated using the cellranger *aggr* pipeline. To ensure only high-quality cells were integrated into further bioinformatic analyses, cells with low total UMI count (< 2500) and high mitochondrial RNA content were filtered using the cellranger *reanalyze* pipeline.

### Single cell RNA-seq pseudotime analysis

Force directed layout (FDL) plots were computed using scanpy, as per the Palantir package for Python^63^. First, aggregated scRNA-seq data (normal tongue (n=2), 5 days (n=1), 2 weeks (n=1), 4 weeks (n=1), 8 weeks (n=2), and 16 weeks (n=3) post TAM) was filtered for minimum cells/gene (>3) and genes/cell (>200), as well as mitochondrial genes (“mt-” were removed). The data was then normalized per cell, log transformed and 3000 variable genes were selected for subsequent principal component analysis, diffusion mapping, FDL computation, magic imputation and visualization. Next, pseudotime trajectories were generated using the *run_palantir* function. Early cells were selected from the centroid of the ground truth normal tongue timepoint cluster. Terminal states were selected from edge cells of two of the 16-week sarcoma samples (clustered together in FDL space). A gene list was manually curated based on differential expression analysis and corresponding gene trends were computed using the *compute_gene_trends* function, with trajectory expression patterns visualized in pseudotime using the *plot_gene_trends* function. For analysis of gene clusters over pseudotime, the data matrix was filtered to remove cycling cells, pericytes, contaminating red blood, and endothelial cells before reanalysis with Palantir. The same start cell and terminal cells were used however the 2 SRC trajectories were collapsed into a single trajectory. Gene clusters were calculated using the *cluster_gene_trends* function on 5000 highly variable genes defined by the harmony *hvg_genes* function and gene cluster trend plots generated using the *plot_gene_trend_clusters* function.

Embryonic fibroblast data was sourced from a previous study^53^, and a list of top differentially expressed genes between E11.5 and E16.5 fibroblasts were compiled by reanalyzing a subset of cells, herein referred to as eMSC to fibroblast program. To obtain fibroblast trajectories, palantir analysis was performed as described above. Gene trend heatmaps, including eMSC and fibroblast developmental program genes and additional manually curated genes of interest from adult fibroblasts and sarcoma cells, were generated using the *plot_gene_trend_heatmaps* function.

### Single cell ATAC-seq

For single cell ATAC-seq captures, ∼100,000 FACS-purified cells were collected from normal tongue, as well as 8- and 16-week (endpoint) tongue tumours, as per dissociation and sorting conditions described above. Nuclei were isolated by lysing the cells for 5 minutes, according to *Nuclei Isolation for Single Cell ATAC Sequencing* (10X Genomics, CG000169 Rev E). Nuclei were quantified using a Countess II FL automated cell counter (ThermoFisher) and ∼10,000 nuclei were targeted for transposition and capture using a 10X Chromium controller. ScATAC-seq libraries were prepared according to the *Chromium Single Cell ATAC Reagent Kits User Guide* (10X Genomics; CG000168 Rev B). Single cell libraries were sequenced on a Nextseq500 or 2000 (Illumina). The *cellranger-atac* pipeline (version 2.1.0) was used to generate fastq files from the sequencer output bcl files, perform read filtering and alignment, and for the detection of accessible chromatin peaks, dimensionality reduction, cell clustering and differential accessibility analyses. Track plots were generated using Loupe Cell Browser (10x Genomics).

### ChIP-Seq

#### Sample preparation

Tissues (normal tongue (n=23) or tongue tumour (n=3)) were dissociated and sorted as described above, with the following differences: tissues were dissociated using activated Collagenase D (1.5 u/mL)/ Dispase II (2.4 u/ mL) digest cocktail only, rotated for 1.5 h at 37°C, and samples were resuspended prior to sorting, and collected post-sort, in PBS containing 2% FBS and 1x protease inhibitor cocktail (PIC Set III, Millipore Sigma, Cat. 539134). For each sample, 200K cells were sorted, counted with a hemocytometer and pelleted by centrifugation at 500 g, with no break, for 6 min at 4°C, after which the supernatant was removed. Cell pellets were resuspended with 50 uL of 0.1% Triton X-100, 0.1% Deoxycholate, 10 mM sodium butyrate and 1x PIC, and immediately flash frozen with liquid nitrogen. ChIP library construction and sequencing were performed following the guidelines formulated by the IHEC, and are available at https://thisisepigenetics.ca/for-scientists/protocols-and-standards. The list of antibodies used is available at https://thisisepigenetics.ca/data/CEMT/metadata/antibody_qc.html. Input DNA libraries were constructed for background correction during peak calling. A customized version of the paired-end sample Prep Kit from Illumina (V.1.1) was used for sample preparation^1^. Libraries were indexed, pooled, and sequenced using paired-end 75 nt sequencing chemistry on an Illumina HiSeq 2500 platform, following the manufacturer’s protocols.

#### ChIP-Seq analyses

The resulting sequence reads were split by index and aligned to the reference mouse genome (mm10) using BWA alignment tool (v0.5.7)^64^. Read-depth normalized BigWig files were generated using deepTools^65^ (v3.3.0) bamCoverage. Model-based analysis of ChIP-seq (MACS2)^66^ was used for peak calling. Read density in protein coding promoters (+- 2kb from the TSS) was calculated using deepTools multiBigwigSummary from the normalized BigWig files. Bedtools^67^ (v2.30.0) was used to calculate the intersect between MACS2 peak calls of H2AK119Ub and other histone marks. In addition, bedtools was used to identify which protein coding promoter regions (+/- 2kb from the TSS) overlapped peak calls of specific histone marks. All data was visualized using ggplot2 (v3.4.4).

ChIP-seq heatmaps were generated using deepTools (version 3.5.4.post1). BigWig files were first processed with computeMatrix in reference-point mode, generating a matrix file containing calculated scores for each gene located upstream and downstream of the TSS site (+/- 5kb). The tool plotHeatmap was utilized to read the matrix files and produce heatmaps.

### Xenium Spatial Transcriptomics

#### Slide Preparation

For Xenium analyses, two slides were prepared; one slide contained five tongue samples (two normal and three tumours) and the second slide consisted of six stifle tumours. Tissue sections were prepared as per the *Xenium In Situ for FFPE – Tissue Preparation Guide* (10X Genomics, CG000578 version Rev C) protocol. Briefly, 5 μm sections were cut, floated on an RNAse-free Milli-Q water tissue flotation bath, and carefully mounted onto the Sample Area of a Xenium slide using a paintbrush. The slides were then air dried in a slide rack at room temperature for 30 min, followed by a 3 h incubation at 42°C in a mechanical convection oven (Precision, Model STM 80). Slides were stored in a sealed container at room temperature with desiccant for three days.

Subsequently, the slides were processed, over a two-day span, as per the *Xenium In Situ for FFPE – Deparaffinization and Decrosslinking* (10X Genomics, CG000580 version Rev C) and *Xenium In Situ Gene Expression* (10X Genomics, CG000582 version Rev C) protocols. In brief, on day one, slides were incubated at 60°C for 2 h on a Xenium thermocycler adaptor plate (BioRad, C1000 Touch Thermocycler), followed by deparaffinized through a series of immersions in xylene, 100%, 96% and 70% ethanol, and rehydration with nuclease-free water. Slides were assembled into Xenium cassettes for all downstream steps, starting with decrosslinking and permeabilization. Gene panel probes were hybridized overnight (minimum 18 h) at 50°C. A total of 348 genes were included in this dataset; 248 from the pre-designed gene expression probe set *Xenium Mouse Brain Gene Expression Panel* (10X Genomics, PN-1999462) and 100 from a custom panel (10X Genomics, XD4MB9). The custom panel was designed based on analysis of single cell data included in this manuscript. On day two, slides were subjected to a post-hybridization wash (37°C for 2 h), ligation (37°C for 2 h), amplification (30°C for 2 h), autofluorescence quenching and nuclei staining steps. Slides were then stored in PBS-T, overnight, at 4°C, in the dark. The following day, slides and consumables were loaded onto the Xenium Analyzer instrument according to the *Xenium Analyzer* (10X Genomics, CG000584, version Rev B) protocol. Regions of interest encompassing entire tongue (normal or containing tumor), or stifle tumor tissue sections were selected for probe decoding image acquisition and the run initiated.

#### Post-run H&E staining

After run completion, slides were removed from the Xenium instrument and stored in PBST for up to 1 hour, until H&E stained as per the “Xenium in Situ Gene Expression – Post Xenium Analyzer H&E Staining” (10X Genomics, CG000613, version Rev A) protocol. Briefly, slides were immersed in quencher removal solution for 10 min, post-fixed in 10% Neutral Buffered Formalin for 5 min, rinsed twice (5 min each) in PBS, and transferred to Milli-Q water prior to standard Hematoxylin and Eosin staining. Slides are stained in hematoxylin (Gill’s II, EMD Millipore, 65066-85) for 1 min and 40 sec, clarified in acid solution (Epredia, 7401) for 30 sec, blued in alkaline solution (in-house formulation, pH=10, sodium bicarbonate and lithium carbonate in 25% (aq) methanol) for 1 min, stained in alcoholic eosin Y (Epredia, 71204) for 28 sec, differentiated and dehydrated in 100% ethanol, cleared in xylenes, and cover slipped with resinous mounting medium (Leica, 3801731).

#### Xenium data analysis

Analysis of Xenium samples and generation of UMAPs, heatmaps and positional gene expression plot figure panels was completed in R (version 4.3.2) using Seurat (version 5.0).

### Quantification and Statistical analysis

All data are represented as mean ± SD, the sample number and statistical analyses are indicated in the figure legends. Sample size determination was based on anticipated variability and effect size that was observed in the investigator’s lab for similar experiments. Kaplan-Meier plots were generated using Prism 8 (GraphPad). For pairwise comparisons, unpaired two-tailed t tests were used to calculate P values. For comparison of >2 means, one-way ANOVA with Bonferroni correction was employed. Both statistical methods were carried out using Prism (GraphPad Software v. 9). p values < 0.05 were considered significant.

## Data availability

The scRNA-seq and scATAC-seq data generated in this study have been deposited to GEO under accession code GSE216769 and GSE213265, respectively. GEO accession code (pending) is associated with RNA-seq and ChIP-seq datasets. The embryonic scRNA-seq dataset from lineage-traced *Hic1^CreERT2^*; *Rosa^tdTom/+^*mice used in Extended Data Fig. 11 is publicly available at GEO repository with accession number GSE156953.

## Code availability

The software used to analyze the data is either freely or commercially available.

## Extended Data Figure Legends

**Extended Data Fig. 1.**
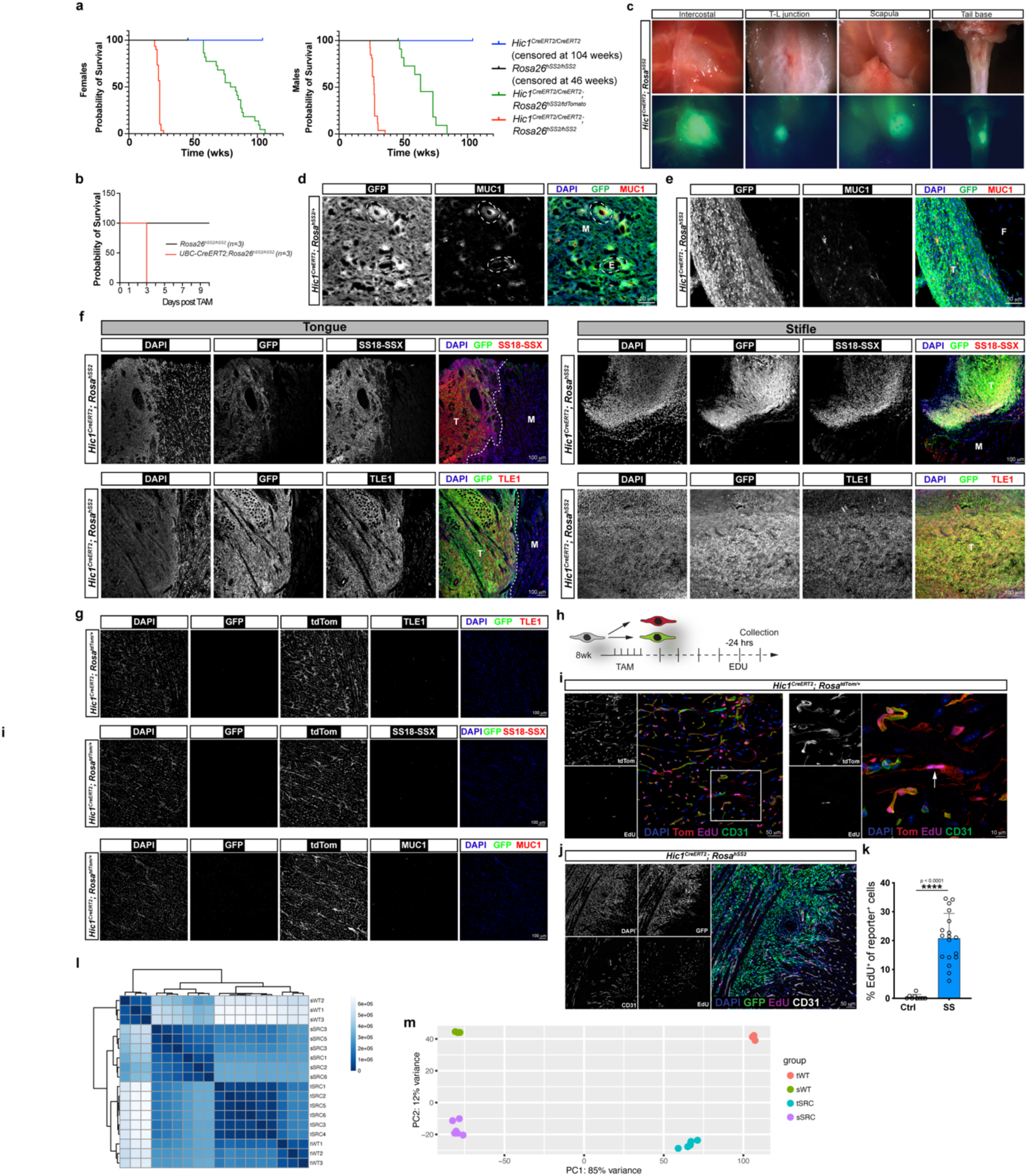
Expression of *SS18::SSX* in *Hic1*^+^ MSCs induces tumours with features of SyS. **a,** Kaplan Meier survival curves for the indicated genotypes, separated based on sex. **b**, Kaplan Meier survival curve for mice in which *SS18::SSX* was ubiquitously expressed using UBC-CreERT2, at 8 weeks of age. **c**, Whole mount images of GFP^+^ tumours (*Hic1^CreERT2^*; *Rosa^hSS2^*) at various anatomical sites. **d-g,** Representative images of immunofluorescence staining of sections from end-point tongue and stifle tumours for MUC1 (**d;** broken lines indicate areas of pseudogland formation), SS18::SSX (**e;** dotted lines demarcate tumour margin) and TLE1 (**f**). T, tumour; F, fascia; M, muscle; E, epithelioid. **g**, Corresponding immunostaining of control tongues (*Hic1^CreERT2^*; *Rosa^tdTomato/+^*). **h**, Schematic overview of the experimental plan for evaluating cell proliferation using EdU incorporation. **i-k**, Detection of EdU^+^ cells in control tongue sections (**i**; White box region is enlarged in right panel with a rare single EdU^+^ tdTom^+^ cell highlighted by arrow) and within GFP^+^ tongue tumours (**j**) collected at end-point (∼16 weeks post-TAM), counterstained with DAPI and the indicated antibodies. **k**, Enumeration of EdU^+^ cells in control and tumour-bearing mice, as a percentage of reporter^+^ cells. **l**, Correlation matrix heatmap of Euclidean distance between control and tumour samples from the tongue and stifle region based on RNA-seq gene expression. **m**, Principal component analysis plot of the variance explained by the first two components across the samples from (l), colored by sample group.

**Extended Data Fig. 2.**
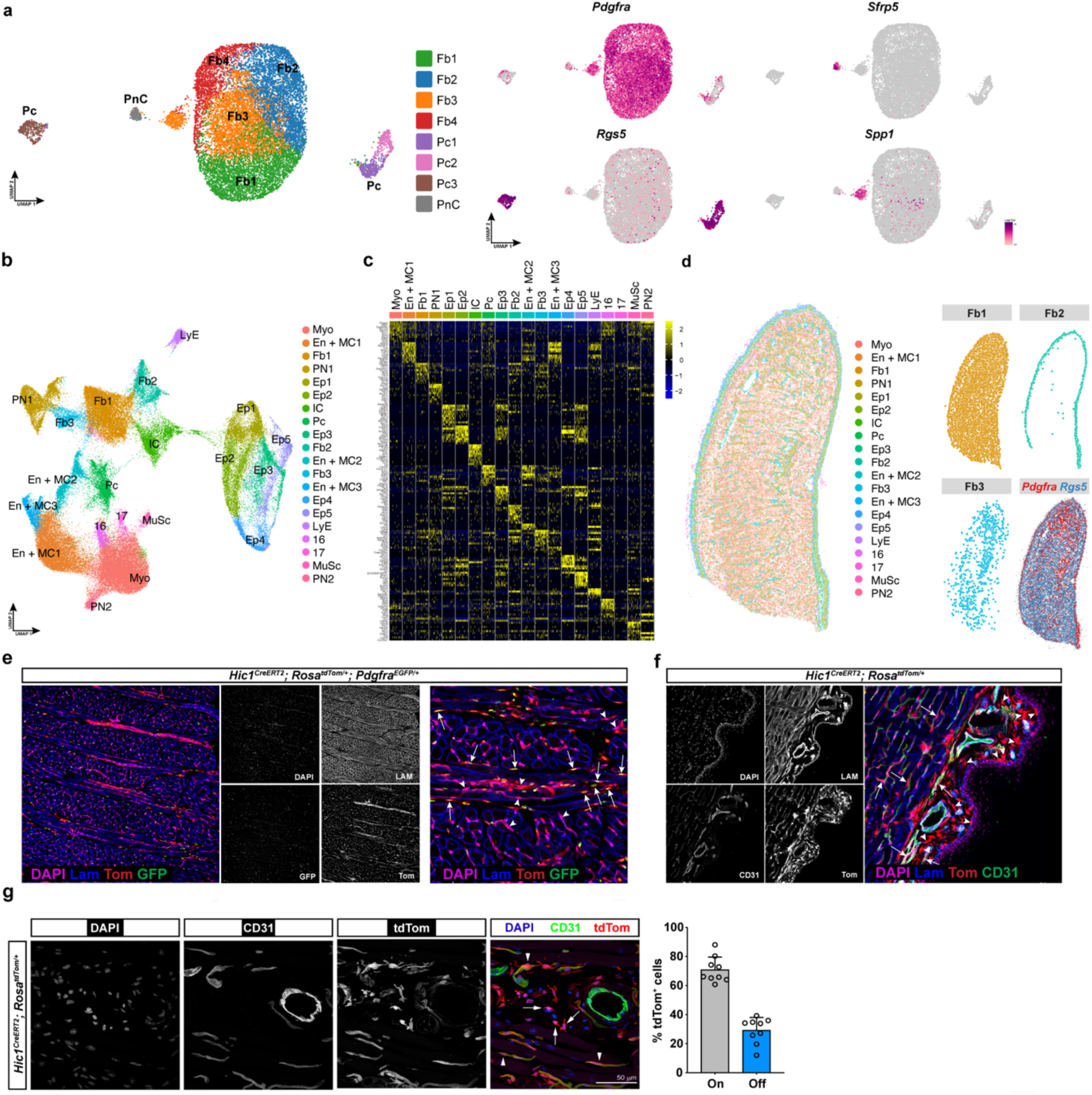
Cellular and molecular characterization of *Hic1*^+^ MSCs in the tongue. **a,** UMAP of scRNA-seq samples (n=2) generated from *Hic1*^+^ MSCs (*Hic1*; tdTom^+^) colored by cluster identity. K-means (k=8) was used for clustering, and cell types are annotated based on transcripts representative of different lineages (right): Fb, fibroblast; Pc, pericyte; PnC, perineural. **b**, UMAP colored by cluster identity derived from spatial transcriptomic analysis, annotated by marker analysis in (c). **c**, Heatmap of top 10 defining transcripts, with their cellular identity defined as: Myo, myofibre; En + MC, endothelial and mural; Fb, fibroblast; PN, peripheral nerves; Ep, epithelial; IC, immune; Pc, pericyte; LyE, lymphatic endothelium; MuSc, muscle satellite cell. **d**, Mapping of identified clusters to spatial domains, with the distribution of Fb cell types highlighted on the right. Spatial distributions of *Pdgfra* and *Rgs5* transcripts are also shown. **e-f,** Representative images of Hic1^CreERT2^; *Rosa^tdTom/+^*; *Pdgfra^EGFP/+^*cells in the medial (e) and ventral (f) compartments of the tongue. Arrows identify tdTom^+^ (Tom) EGFP^+^ (GFP) cells; arrowheads denote Tom^+^ GFP^-^ cells. Nuclei, basement membrane and vasculature counterstained with DAPI, anti-Laminin and anti-CD31, respectively. **g**, Representative images of *Hic1^CreERT2^*; *Rosa^tdTom/+^* cells within the tongue, and enumeration of their proximity to CD31-positive vessels (n=3, 3 independent sections from 3 biological replicates).

**Extended Data Fig. 3.**
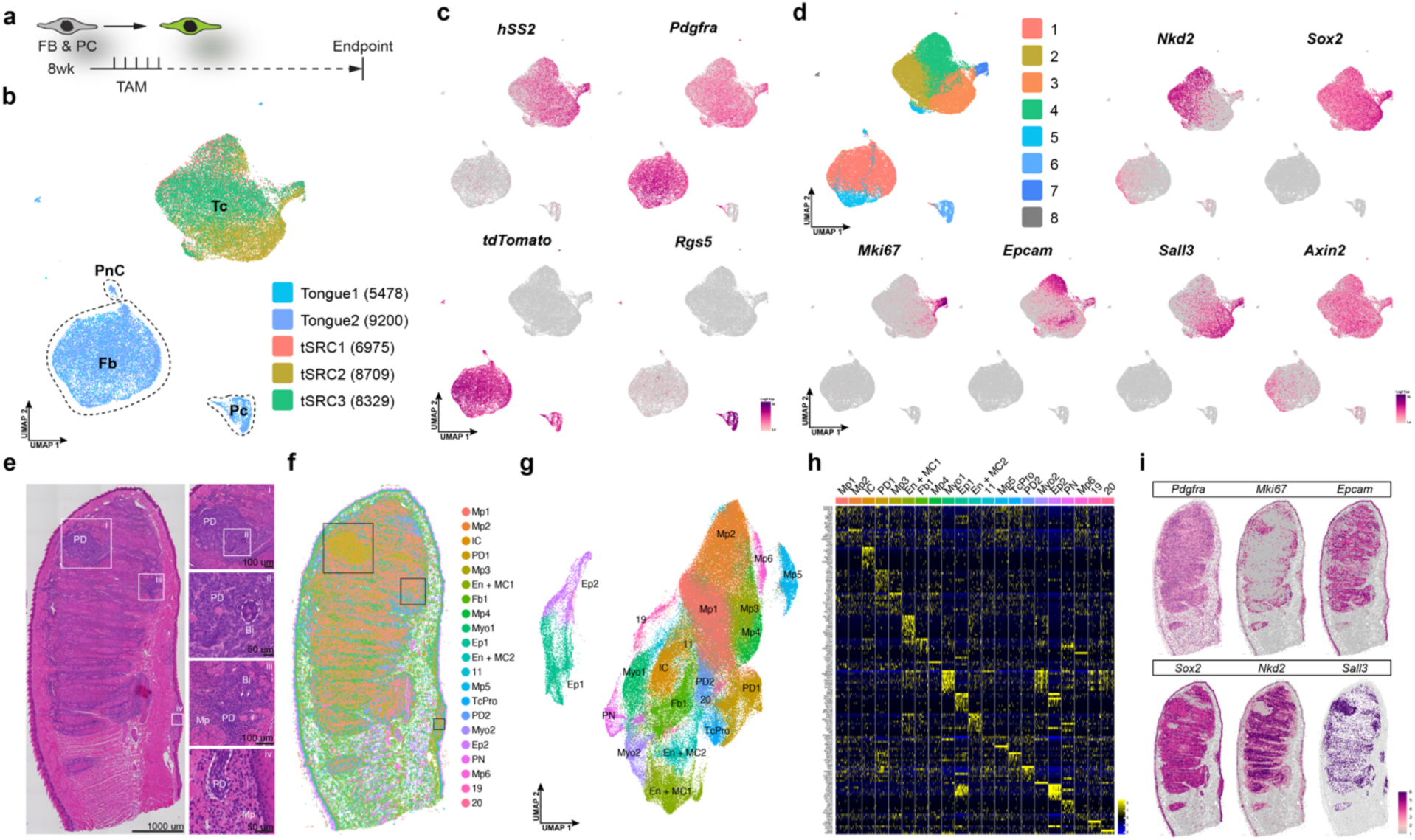
Cellular and spatial transcriptomic profiling of tongue tumours. **a**, Schematic overview of experimental plan. **b**, ScRNA-seq UMAP of cells collected from 3 end-point tongue tumours (tSRC1-3) and two wild-type controls (Tongue1 and 2) colored by sample ID. Fb, fibroblast; Pc, pericyte; PnC, perineural cells; and Tc, tumour cell. Wild-type control cells were derived from *Hic1*; *tdTom*^+^ mice (10 days post-TAM). **c**, UMAPs of scRNA-seq colored by gene expression level of select indicated transcripts. **d**, UMAP from (b) colored by cluster identity (K-means, k=8) of the, with cluster-enriched transcripts shown. **e**, H&E whole-mount section of a representative tongue tumour (tSRC2) processed for spatial transcriptomics, with insets (i, ii, iii, iv). Histological sub-types are labelled; Bi, biphasic; Mp, monophasic; PD, poorly differentiated. **f**, Cell clusters identified based on transcript expression, mapped to spatial domains on the image in (e). Boxed regions correspond to insets in (e). **g**, UMAP of cells identified in (f) colored by cluster identity, with their assignment to different tumour phenotypes and cell types specified: IC, immune; Myo, myofibre; En + MC, endothelial and mural; Fb, fibroblast; Ep, epithelial. **h**, Heatmap of the top 10 cluster markers for the clusters in (f-g). **i**, Spatial localization of select transcripts colored by expression level mapped to images in (e-f).

**Extended Data Fig. 4.**
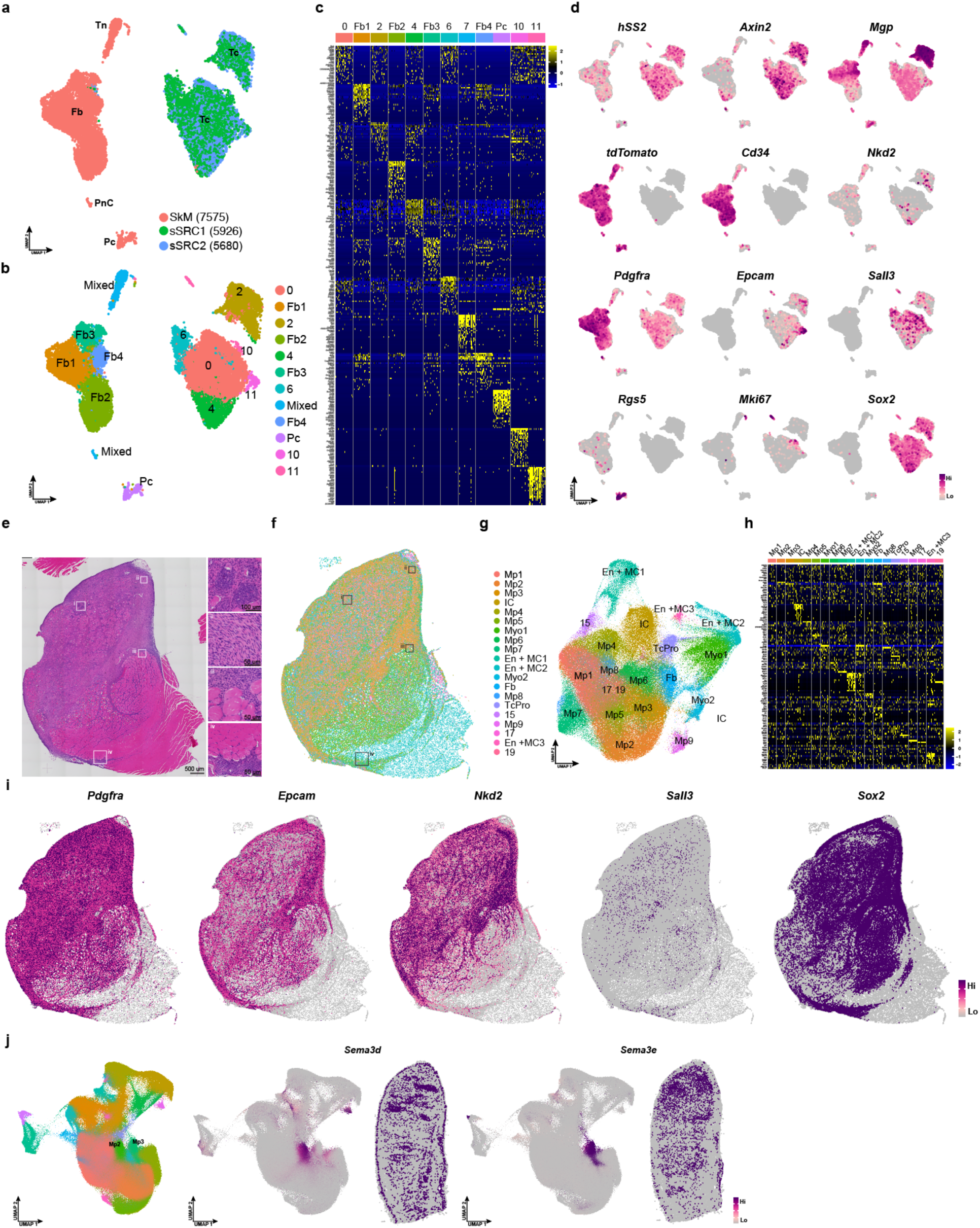
Cellular and spatial transcriptomic profiling of stifle and tongue tumours. **a-b,** UMAPs of scRNA-seq data aggregated from the indicated samples, colored by sample ID (a) and cluster ID (b). SkM, hind limb skeletal muscle with overlying facia; sSRC, stifle sarcomas. Cell types are also indicated: PnC, perineural cell; Pc, pericyte; Tc, tumour; Tn, *Scx*^+^ tenogenic. **c**, Heatmap of scRNA-seq data showing the top 25 enriched genes in the 12 different clusters in (b). **d**, UMAPs from (a-b), colored by expression level of various cluster-enriched transcripts. **e**, H&E whole-mount section of a representative stifle tumour (sSRC3) processed for spatial transcriptomics, with insets (i, ii, iii, iv). **f**, Cell clusters identified based on transcript expression, mapped to spatial domains on the image in (e). **g**, UMAP of cells identified in (f) colored by cluster identity, with their assignment to different tumour phenotypes and cell types specified; IC, immune; Myo, myofibre; En + MC, endothelial and mural; Fb, fibroblast. **h**, Heatmap of the top 10 cluster markers for the clusters in (f-g). **i**, Spatial localization of select indicated transcripts in stifle tumours colored by expression level. **j**, UMAPs and spatial localization of select transcripts in tongue tumours colored by expression level mapped to images in (Ext data 3e-f).

**Extended Data Fig. 5.**
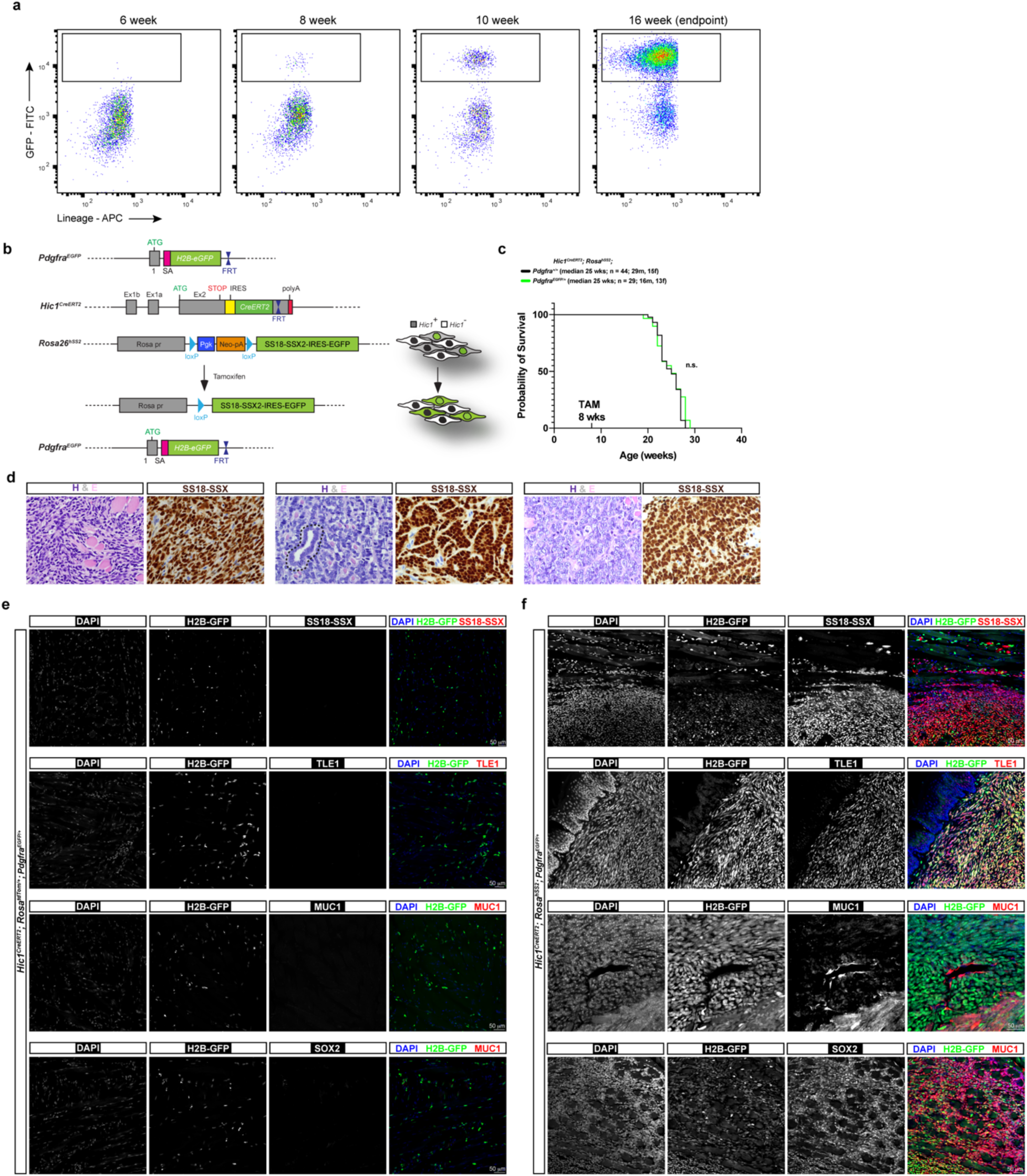
Haploinsufficiency of *Pdgfra* does not affect synovial sarcomagenesis. **a,** Flow cytometry analysis of GFP^+^ cells collected at the indicated time points from *Hic1*; *hSS2* tumour samples. **b,** Schematic of the genetic strategy employed to incorporate a *Pdgfra^EGFP/+^* allele into the *Hic1*; *hSS2* SyS model. **c**, Kaplan Meier survival plot of the indicated genotypes treated with TAM at 8 weeks. n.s., not significant. **d**, H&E and anti-SS18::SSX staining of end-point tongue tumours from the *Hic1^CreERT2^*; *Rosa^hSS2/hSS2^*; *Pdgfra^EGFP/+^* background, left to right, examples of monophasic, biphasic with pseudoglandular structure (dotted line) and poorly differentiated. **e**-**f**, Images of tongue control (**e**, *Hic1^CreERT2^*; *Rosa^tdTom/+^*; *Pdgfra^EGFP/+^*) and end-point tumour (**f**, *Hic1^CreERT2^*; *Rosa^hSS2/hSS2^*; *Pdgfra^EGFP/+^*) samples immunostained for the indicated targets.

**Extended Data Fig. 6.**
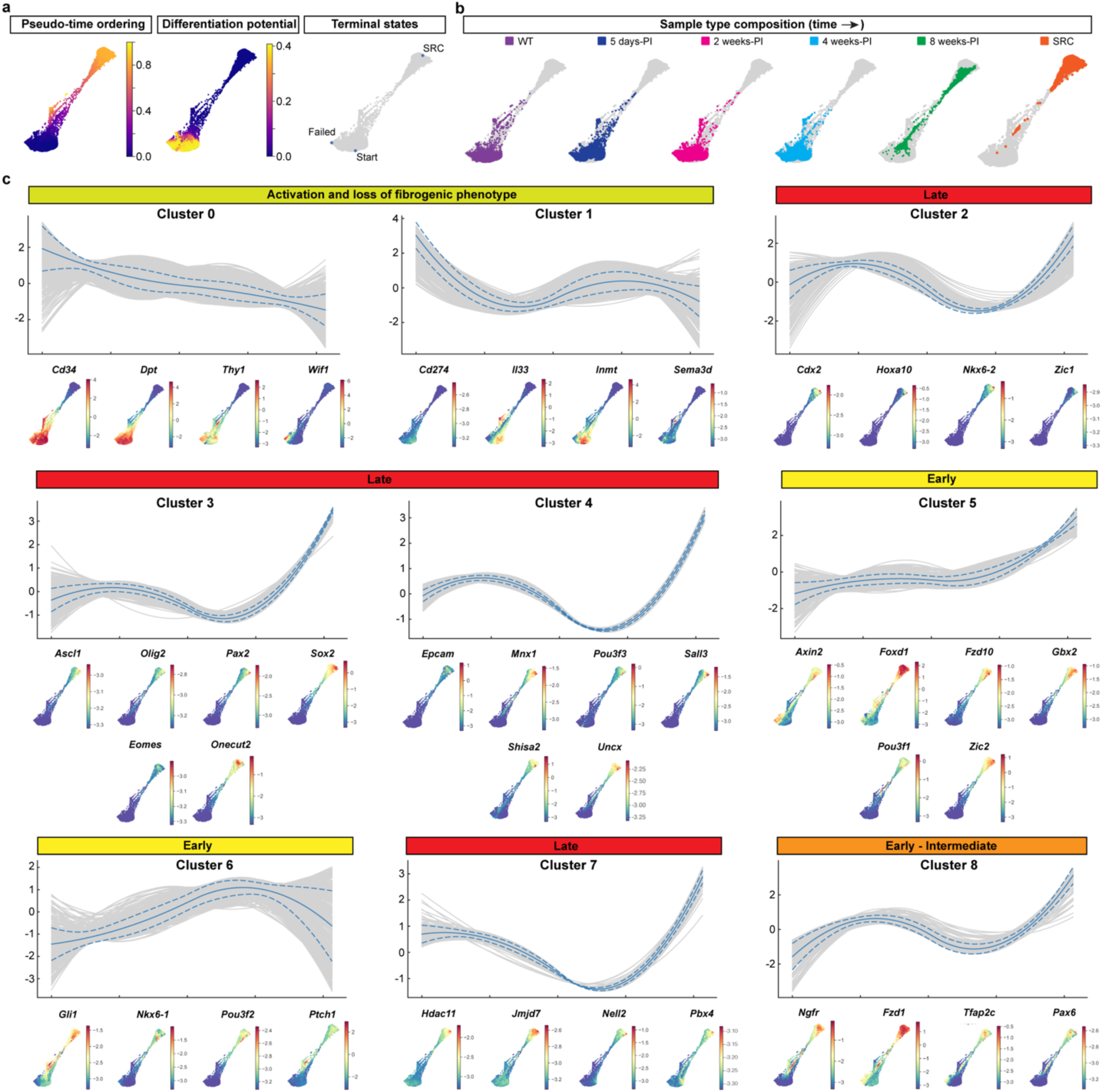
Delineation of the temporal transcriptional program underlying SS18::SSX-mediated reprogramming. **a**, Force-directed layout (FDL) plot of filtered, aggregated data (from Fig. 3d-h) colored by pseudo-time ordering of cells (left), predicted differentiation potential (middle) and the cells selected for trajectory terminal points used for further analysis (right). **b**, FDL plots from (a-b), highlighting in color the indicated actual time points of sample collection within each plot. **c**, Gene trend plots of calculated gene clusters along the sarcomagenic trajectory defining the indicated stages of the transformation process. FDL plots colored by the expression levels of representative genes from each cluster (below).

**Extended Data Fig. 7.**
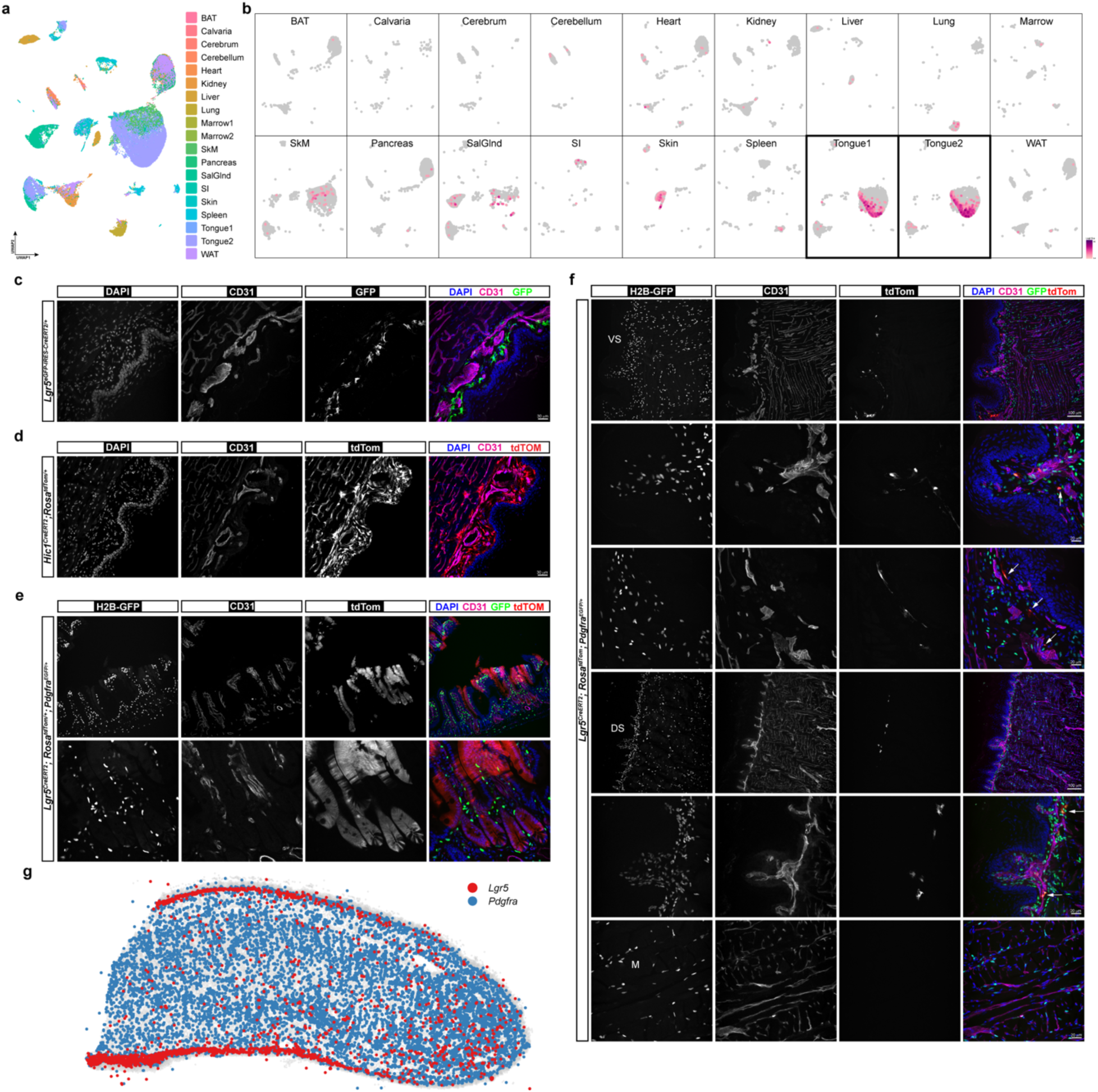
Characterization of *Lgr5*^+^ fibroblasts in the tongue. **a,** UMAP of aggregated scRNA-seq data colored by sample ID comprising *Hic1*; *tdTom*^+^ MSCs, isolated and enriched from the indicated tissues of 10-week old mice, 10 days post-TAM induction. BAT, brown adipose tissue; Marrow, bone marrow from fore and hindlimb bones; SalGlnd, salivary gland; SI, small intestine; WAT, white adipose tissue. **b**, UMAP from (a), split by tissue type, colored by *Lgr5* expression level. **c-f**, Representative images of: **c**, Immunodetection of GFP and CD31 in the ventral lamina propria of *Lgr5^CreERT2-GFP^* mice. **d**, Detection of the tdTom^+^ population (*Hic1^CreERT2^*; *Rosa^tdTom/+^* mice) within the ventral region of the tongue. **e**, Lineage tracing of *Lgr5* progeny in the small intestine using the Rosa^tdTom*/+*^ reporter gene in the indicated genetic background. Samples were collected 10 days post-TAM administration. **f**, Detection of *Lgr5* lineage traced cells (tdTom^+^) and *Pdgfra*^+^ cells (nuclear H2B-GFP*^+^*). Arrows denote doubly positive tdTom and nuclear GFP cells in the dorsal and ventral regions of the tongue. DS, dorsal surface; M, midline; VS, ventral surface. **g**, Distribution of *Lgr5* and *Pdgfra* transcripts in the wild-type tongue, as determined by spatial transcriptomic analysis.

**Extended Data Fig. 8.**
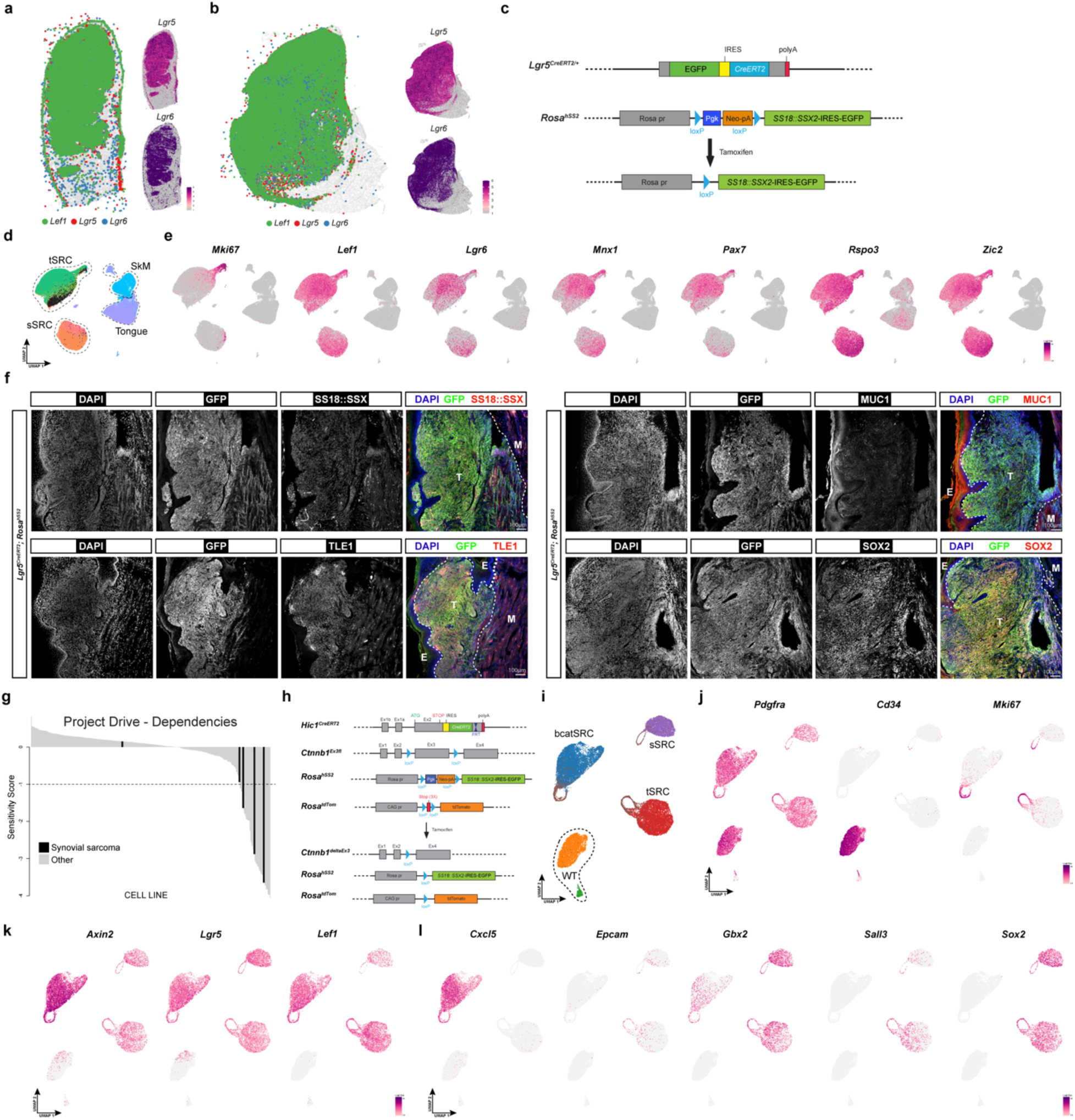
Beta-catenin/WNT signaling in synovial sarcomagenesis. **a-b,** Spatial transcriptomic analysis of *Lef1*, *Lgr5* and *Lgr6* distribution in tongue (**a**) and stifle (**b**) tumours. **c**, Experimental plan for induction of *SS18::SSX* in *Lgr5*-expressing cells. **d**, scRNA-seq data UMAP of cells from the indicated samples colored by sample ID with the *Lgr5*-tumour derived sample highlighted in black. **e**, UMAPs from (d) colored by the expression level of the indicated select transcripts. **f**, Representative images of the immunodetection of tumour-associated proteins (SS18::SSX, TLE1, MUC1 and SOX2), in relationship to GFP^+^ tumour cells, from sections of *Lgr5^CreERT2^; Rosa^hSS2/hSS2^* tumours, counterstained as indicated. **g**, Waterfall plot of *CTNNB1* dependencies as determined in the DRIVE project. Each black line indicates a different human synovial sarcoma cell line. **h**, Schematic of the alleles used to activate beta-catenin/WNT signaling in the *Hic1*; *hSS2* background. **i-l**, UMAPs of scRNA-seq data, colored by sample ID (**i**) and by the expression level of select differentially expressed genes representative of normal fibroblasts (**j**), WNT signalling components (**k**) and mid-to-late stage transformed cells (**l**).

**Extended Data Fig. 9.**
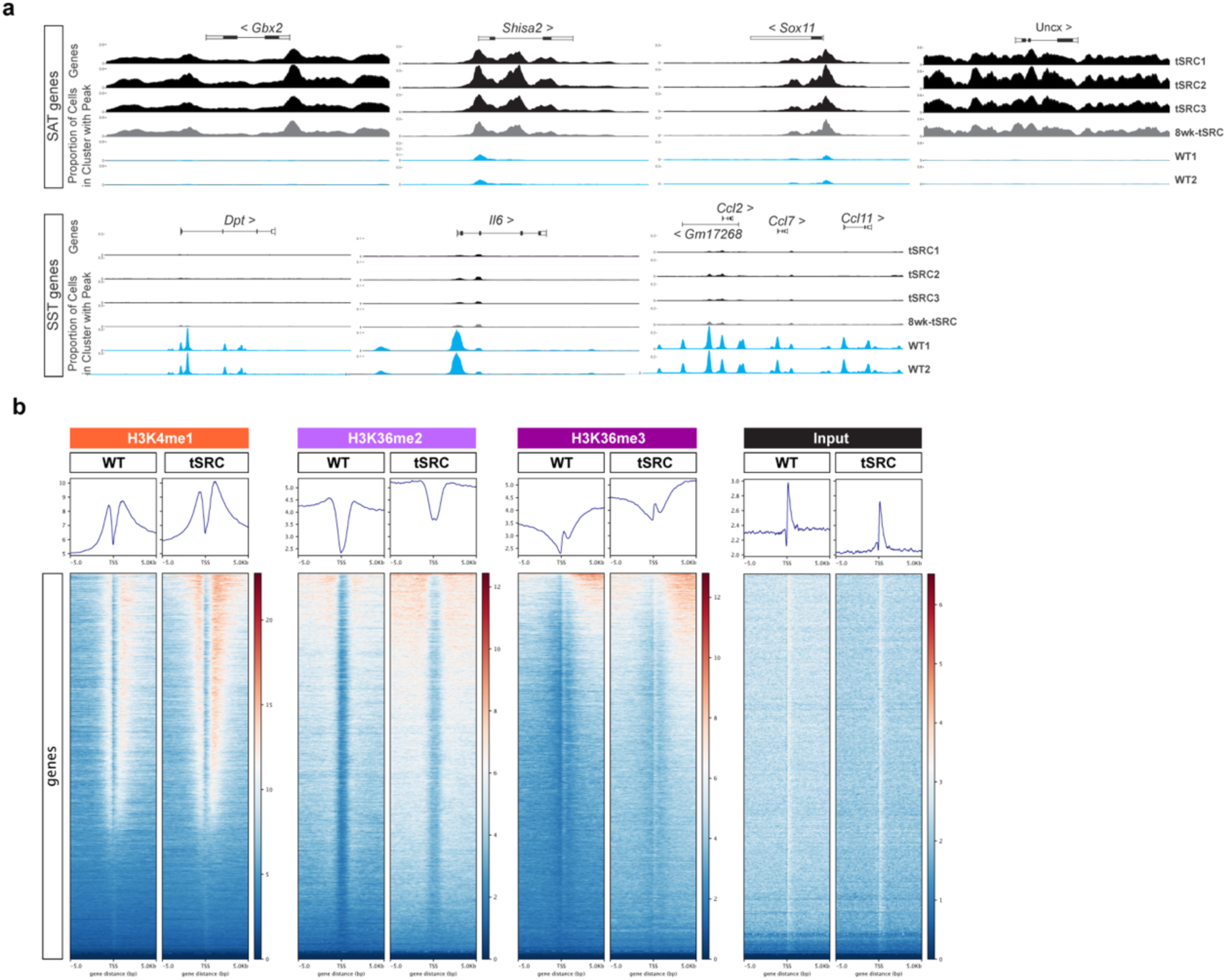
Modification of the epigenome by SS18::SSX. **a,** Genome browser tracks derived from scATAC-seq, displaying the promoter sum accessibility signal around specified gene loci from the indicated samples. tSRC, endpoint (∼16 weeks post TAM) tongue tumour; 8wk-tSRC, 8 weeks (post TAM) tongue tumour; WT, wild-type. **b**, Profile heatmaps of the indicated histone modifications, detected by ChIP-seq, around the TSS (+/- 5kB) of RefSeq genes. The gradient, blue-to-red, represents low-to-high counts in the noted regions.

**Extended Data Fig. 10.**
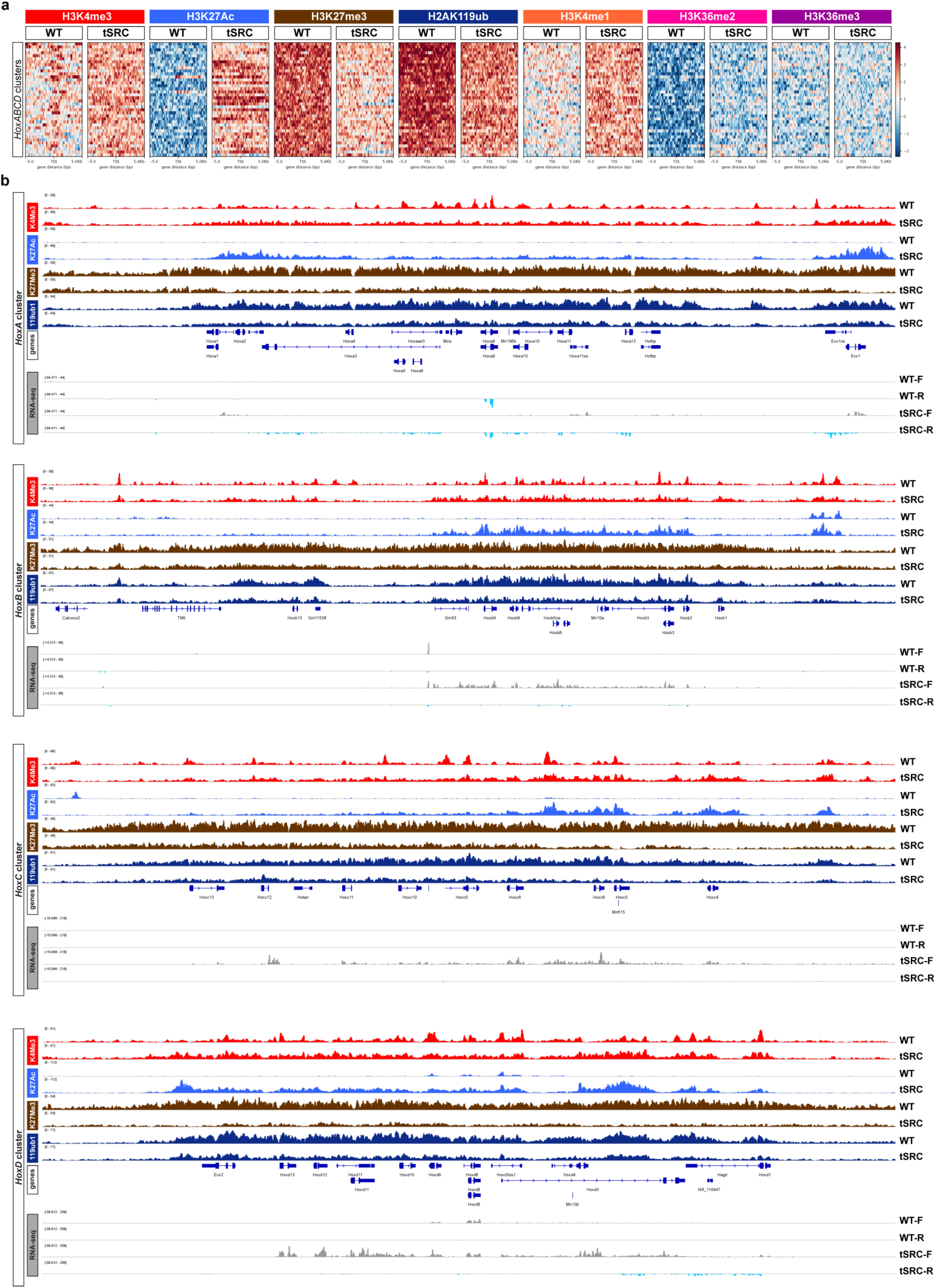
Broad “re-activation” of genes within the HoxA, B, C and D clusters through SS18::SSX-mediated reprogramming. **a**, Heatmaps of indicated histone modifications observed at the *HoxA*, *B*, *C* and *D* loci (TSS +/- 5kB). **b**, Individual genome track plots of indicated histone modification peaks detected by ChIP-seq, along with transcript abundance generated from RNA-seq (+ strand (F) and – strand (R) tracks), within the HoxA, B, C and D clusters. Annotation track denotes exon and intron structure, along with coding information. WT, wild-type; tSRC, endpoint tongue tumour.

**Extended Data Fig. 11.**
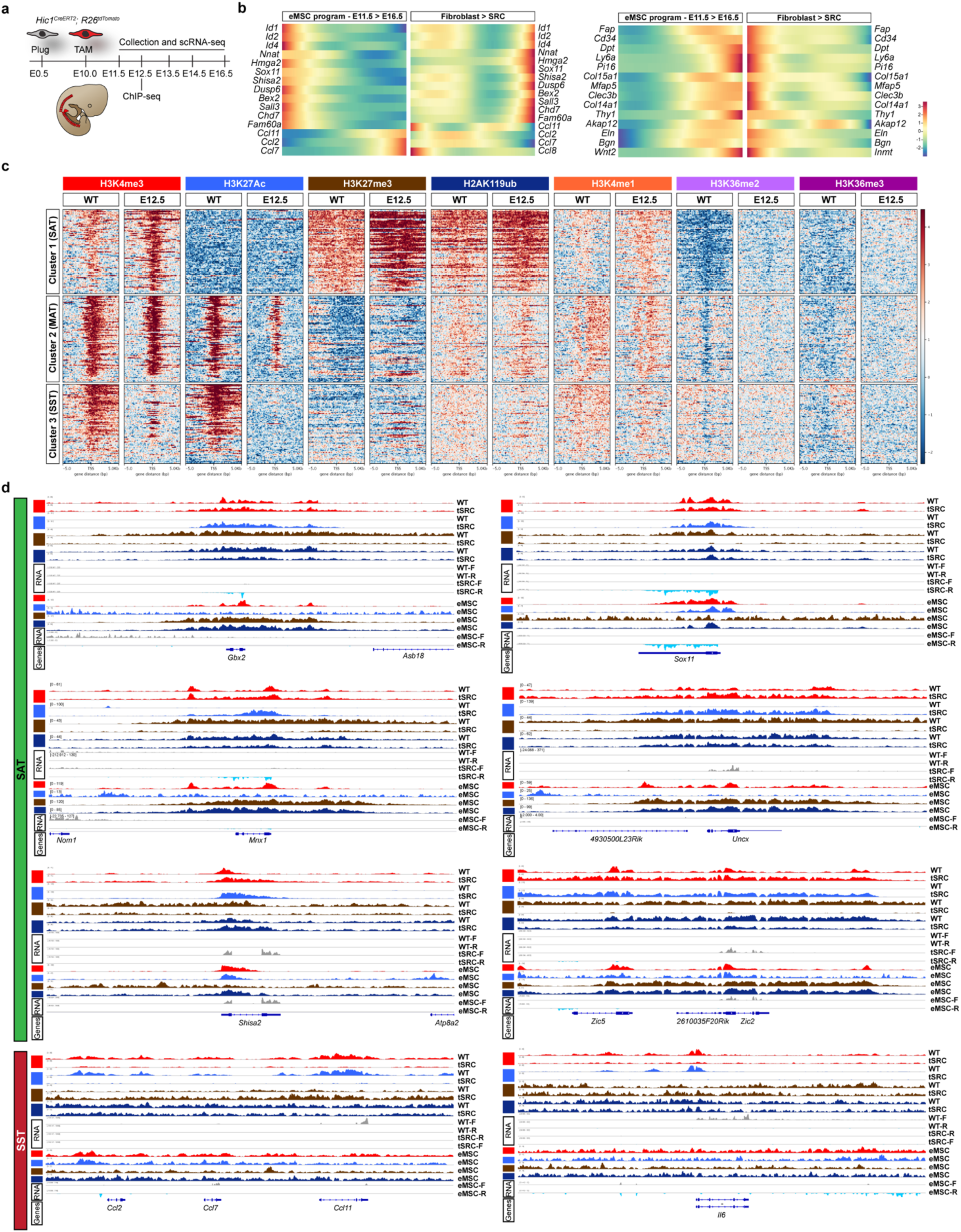
Adult fibroblasts harbour an epigenome reflective of their embryonic origin. **a,** Overview of the experimental paradigm used for collection of omics data from embryonic MSCs (eMSCs). The red banding pattern indicates where *Hic1*^+^ cells are first detected in the somitic compartment of the embryo^53^. **b,** Pseudotime trajectory based heatmaps of select genes in *Hic1*; *tdTom*^+^ MSCs that exhibit differential expression as they mature during embryonic limb development (E11.5 to E16.5), along with corresponding trajectory heatmaps of fibroblast transformation to SRC (*Hic1*; *hSS2*) cell trajectories. **c**, Heatmaps of indicated histone modifications detected by ChIP-seq for select sarcomagenesis activated (SAT), maintenance (MAT) and sarcomagenesis silenced (SST) transcripts, from 75 loci (TSS +/- 5kB) per cluster. **d**, Individual genome track plots of indicated histone modification peaks detected by ChIP-seq at specific SAT and SST loci, along with transcript abundance generated from RNA-seq (+ strand (F) and – strand (R) of tdTom^+^ enriched wild-type (*Hic1*; *tdTom*) embryonic (E12.5), adult MSCs, and GFP^+^ enriched tumour (*Hic1*; *hSS2*) cells. Annotation track denotes exon and intron structure, along with coding information.

